# Formation of stress granules and non-canonical survival responses in arsenite-exposed cells

**DOI:** 10.1101/2024.07.29.605725

**Authors:** Seishiro Hirano, Osamu Udagawa, Sanae Kanno

**Author notes:** Correspondence author: Seishiro Hirano.

## Abstract

The concentration-dependent decrease in viable cells is a well-documented phenomenon in cytotoxicity assays for most toxic substances. We report that arsenite (As^3+^), a widely recognized oxidative toxicant, exhibited lower cytotoxic effects at 300 µM compared to 100 µM As^3+^ in CHO-K1 and Jurkat cells. Formation of stress granules (SGs), which appear in the cytoplasm shortly after exposure to hypertonicity, heat shock, and high concentrations of As^3+^ is considered as a pro-survival cellular event. We hypothesized that unusual cytotoxicity profile of As^3+^ could be attributed to SG formation. In both CHO-K1 and Jurkat cells stably expressing GFP-tagged G3BP1, SGs were more rapidly and distinctly induced by 300 µM As^3+^ than 100 µM As^3+^. Other toxic metals and a metalloid such as Cd^2+^, Cu^2+^, Ag^+^, and Se^4+^ did not clearly induce SG formation and instead reduced the viability in a concentration-dependent manner. Exposure to As^3+^ led to phosphorylation of eIF2α, a key regulator of polysome stability and a hallmark of SG formation. Depletion of intracellular glutathione (GSH) increased the susceptibility of cells to As^3+^, highlighting its role in cellular defense mechanisms. Exposure to As^3+^ activated small ubiquitin-like modifier (SUMO) which is implicated in phase separation. However, neither depletion of GSH nor overexpression of SUMO contributed As^3+^-induced SG formation. Consistently, THP-1 and HL60 cells exposed to As^3+^ also exhibited non-canonical cytotoxic features, albeit at higher concentrations (1 mM). These findings underscore the need for further mechanistic investigations into As^3+^-induced SG formation, given that As^3+^ is a promising anti-cancer agent, and resistance of tumor cells to As^3+^ is a critical issue.

## Introduction

Arsenic, an ubiquitous environmental contaminants, is well-known for its dual role as a carcinogen [1, 2] and an effective therapeutic agent for acute promyelocytic leukemia [3] and multiple myeloma [4]. Trivalent arsenic (arsenite, As^3+^) is absorbed rapidly by cells and interacts with cysteine residues including those in glutathione [5], leading to the widely accepted notion that As^3+^ induces oxidative stress in living systems.

Other toxic elements such as Cd^2+^ and Hg^2+^ are also known to bind to cysteine and cysteine residues of proteins [6, 7]. However, cellular proteins do not uniformly respond to these thiol-reactive toxicants. Exposure to As^3+^ alters the solubility of promyelocytic leukemia (PML) proteins and induces SUMOylation of the proteins [8, 9]. These biochemical changes are caused by As^3+^ or antimony ions (Sb^3+^), but no other metals and metalloids have been reported to modulate PML proteins [8, 10]. As^3+^ and selenium ions (Se^4+^) have been reported to induce SGs [11], a phenomenon not clearly observed with other metals and metalloids. As^3+^ induces canonical eIF3-containing SGs by eIF2 phosphorylation-dependent mechanism, akin to heat and hyperosmotic stress, whereas SGs induced by Se^4+^ do not contain eIF3 [11–14].

It has been demonstrated that As^3+^ exhibited a biphasic cytotoxic effect in HeLa cells and Gle1, a regulator of DEAD-box family proteins, was essential for SG formation and survival of cells at higher As^3+^ concentrations [15]. SG formation in the cytoplasm promote cell survival by recruiting and protecting mRNAs from installed polysomes [16]. Recent research suggests that SGs also form at sites of endolysosomal membrane rupture, contributing membrane repair [17]. Moreover, SGs enable cells to survive various stresses, including As^3+^ exposure, by inhibiting NLRP3 inflammasome activation and subsequent pyroptosis [18]. Unlike oxidative stressors such as hydrogen peroxide and X-ray irradiation, which induce apoptosis without SG formation, heat shock and As^3+^ generate reactive oxygen species and SGs in cells [19].

In the present study we report that As^3+^-induced cytotoxicity exhibits a non-canonical biphasic curve, and viability at 300 µM was higher than that at 100 µM. This non-canonical cytotoxicity is attributed to SG formation. Interestingly, neither GSH nor SUMO molecules, known determinants of As^3+^-induced cellular events, appear to contribute directly to SG formation.

## Materials and Methods

### Chemicals

Sodium *m*-arsenite (NaAsO_2_), sodium selenite (Na_2_SeO_3_), and phenylarsine oxide (PAO) were obtained from Sigma-Aldrich (St Louis, MO). Cadmium chloride hemi pentahydrate (CdCl_2_ꞏ2.5H_2_O), copper (II) sulfate pentahydrate (CuSO_4_ꞏ5H_2_O), silver nitrate (AgNO_3_), antimony (III) chloride (SbCl_3_), 5-sulfosalicylic acid (SSA), L-buthionine-(S,R)-sulfoximine (BSO), and (+/-)-dithiothreitol (DTT) were purchased from Fujifilm-Wako (Osaka, Japan) and GSH assay kit from Dojindo (Kumamoto, Japan). Necrostatin-1 (necroptosis inhibitor) and ML792 (SUMO E1 inhibitor) were purchased from Cayman (Ann Arbor, MI) and MedKoo Bioscience (Morrisville, NC), respectively. Z-VAD-FMK (pan-caspase inhibitor), Ac-DEVD-CHO (caspase-3 inhibitor), and CellTiter 96 Aqueous MTS Reagent were purchased from Promega (Madison, WI). Lipofectamin LTX and alamarBlue^®^ were purchased from Invitrogen-ThermoFisher (Carlsbad, CA) and BCA protein assay kit from Pierce-ThermoFisher (Rockford, IL).

### Cells

THP-1 (EC 88081201) and Jurkat E6.1 (EC 88042803) cells were purchased from DS PharmaBiomedical (Osaka, Japan). CHO-K1 (RCB 0285), HEK293 (RCB 1637), and HL60 (RCB 041) cells were obtained from RIKEN (Ibaraki, Japan). CHO-K1, Jurkat, and HEK293 cells were transfected with human GFP-tagged human G3BP1 plasmid and stable transfectants selected by neomycin were designated as CHOG3BP1, JurG3BP1, and HEKG3BP1, respectively. CHOG3BP1 cells were further transfected with mCherry-SUMO2 plasmids and stable transfectants expressing wild-type or C-terminal GG-deleted SUMO2 were selected by puromycin. See ‘Source table’ for plasmid information. Jurkat, THP-1, and HL60 cells were cultured in RPMI1640 containing 100 units/mL penicillin, 100 μg/mL streptomycin, and 10% heat-inactivated fetal bovine serum (FBS). CHO and HEK cells were cultured in F12 and DMEM complete culture medium, respectively.

### Measurement of total GSH

CHO cells were culture in a 24-well culture dish, and the cell monolayer was washed once with PBS. The suspension of Jurkat cells were centrifuged and the pellet was washed once in PBS. The cells were lysed by freezing and thawing 2 times in 0.01N HCl solution. An aliquot was used for determination of protein concentrations using a BCA protein assay kit. SSA was added to the cell lysate at a final concentration of 1% and the sample was centrifuged at 8,000 g for 10 min to obtain clear protein-free supernatants. An aliquot of the supernatant was used for total GSH measurement using a GSH assay kit.

### Cell viability

Unless otherwise mentioned, the cell viability was measured using MTS reagent. Briefly, the cells were pre-cultured and exposed to As^3+^ in a 96-well culture dish. After exposure, MTS solution was added to each well and O.D. at 485 nm was measured using a microplate reader (POLARstar OPTIMA, BMG Labtech, Offenburg, Germany). Alternatively, the viability of the cells was assayed by alamarBlue reagent using a fluorescence mode. The cytotoxicity of DTT in Jurkat cells was evaluated by counting viable and dead cells differentially using a traditional trypan blue dye exclusion method with a hemacytometer.

### Western blot

Cells were rinsed with PBS and lysed with ice-cold RIPA buffer containing protease inhibitors (Santa Cruz, Dallas, TX) and phosphatase inhibitor cocktails (Calbiochem-Merck, San Diego, CA) on ice for 10 min. The cell lysate was centrifuged at 9,000 g for 5 min at 4 °C and the supernatant was collected. The protein concentration of the supernatant was adjusted at 2.5 mg/mL. The cellular pellet was rinsed with PBS and digested in Tris-HCl buffer containing 240 U/mL Benzonase^®^ (Santa Cruz) at 25 °C for 2 h with intermittent mixing (Thermomixer comfort, Eppendorf, Wesseling-Berzdorf, Germany). The volume of Tris-HCl buffer was adjusted so that volumes of the RIPA-soluble and -insoluble fractions were the same. Proteins were resolved by SDS-PAGE and then electroblotted onto a PVDF membrane. The membranes were blocked and reacted with primary and secondary antibodies sequentially. Membranes were soaked in ECL (Prime, GE Healthcare, Buckinghamshire, UK) solution, and chemiluminescent images were captured by a CCD camera (ImageQuant 800, GE Healthcare, Uppsala, Sweden). See ‘Source table’ for the primary and secondary antibody information. The membrane was stripped in warmed (56 °C) Tris-HCl buffer (62.5 mM, pH 6.7) containing 100 mM 2-mercaptoethanol and 2% SDS with vigorous shaking for 2 min for reprobing.

### Fluorescent microscopy

CHOG3BP1 cells were cultured in an 8-well cover glass bottom chamber (Iwaki-AGC, Tokyo, Japan) and were treated with As^3+^ or sorbitol. JurG3BP1 cells were treated with As^3+^ or sorbitol in a suspension culture using standard culture dish. For heat treatment, an aliquot of JurG3BP1 cell suspension was transferred in a microcentrifuge tube and the tube was placed in 42.5 ℃ water. After the treatment JurG3BP1 cells were transferred into the cover glass bottomed chamber for microscopic observation. The formation of SGs was captured as live images by fluorescent microscopy (BZ-X710, Keyence, Osaka, Japan, or Eclipse TS100, Nikon, Tokyo, Japan).

### Statistical analyses

Statistical analyses were performed by analysis of variance (ANOVA) followed by Tukey’s *post-hoc* comparison. Statistical significance was accepted when a probability value was less than 0.05. Non-linear curves were fit to the cell survival data when applicable (concentration-dependent) to calculate EC_50_ values using GraphPad prism 5 (GraphPad Software, San Diego, CA)

## Results

### Cytotoxicity of As^3+^ in CHO cells

CHOG3BP1 and CHO-K1 cells were exposed to pre-determined concentrations of As^3+^ for 24 h, and the cell viability was assayed with or without washing the cell monolayer with PBS (Fig. 1). The cell monolayer is commonly washed before the viability assay using adherent cells to remove detached cells, cell debris, and cellular contents released from damaged cells which may interfere with the colorimetric assay. Following washing, viability appeared to be decreased by As^3+^ in a concentration-dependent manner with calculated EC_50_ values of 67 µM for CHOG3BP1 and 73 µM for CHO-K1 cells (Fig. 1A and C). However, when the assay was performed without washing, the viability at 300 µM As^3+^ was unexpectedly higher than at 100 µM As^3+^ in both cell types (Fig. 1B and D). These unexpected results of As^3+^-induced changes in viability were visually confirmed (Supplementary Fig. 1). CHOG3BP1 cells were exposed to pre-determined concentrations of Cd^2+^, Cu^2+^, Ag^+^, and Se^4+^ to investigate whether non-canonical cytotoxicity was specific to As^3+^. Cells exposed to these metal and metalloid ions exhibited typical survival curves or normal concentration-dependent responses (Supplemental Fig. 2), suggesting that the reversed cytotoxicity observed at 300 µM was specific to As^3+^.

**Fig. 1.**
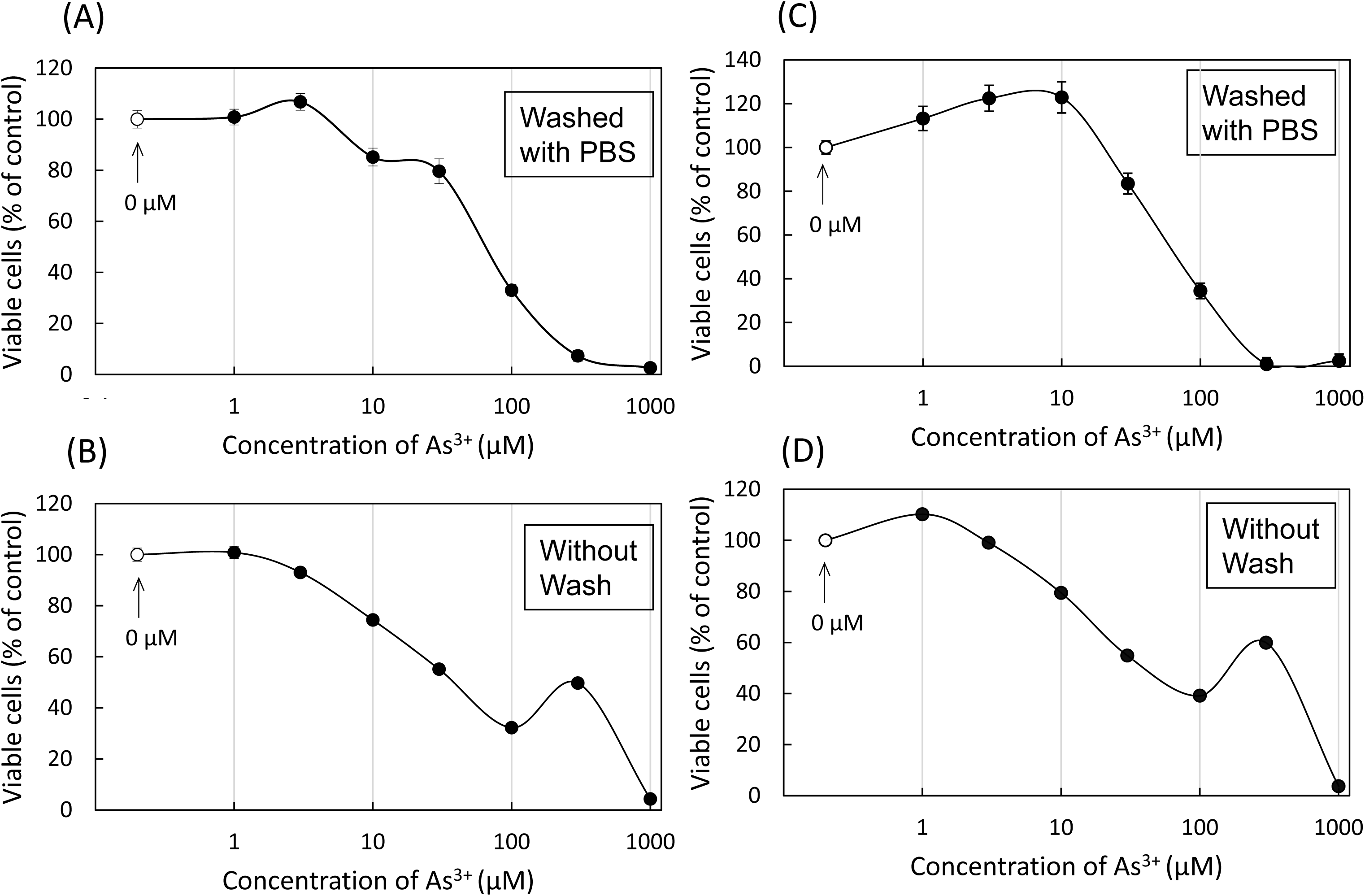
Cytotoxicity of As^3+^ in CHOG3BP1 and CHO-K1 cells. CHOG3BP1 cells (A and B) and CHO-K1 cells (C and D) were exposed to pre-determined concentrations of As^3+^ for 24 h. After exposure, cell monolayers were washed twice with Ca^2+^- and Mg^2+^-free PBS (A and C) or left unwashed (B and D). The cell viability was assayed using the MTS reagent. Data are presented as means ± SEM (N=4).

### SG formation in CHOG3BP1 cells

We hypothesized that the reversal As^3+^ cytotoxicity at 300 µM might be due to SG formation which is known to be pro-survival under lethal stress conditions. CHOG3BP1 cells were exposed to 100 and 300 µM As^3+^ for 0.5 and 1 h and SG-positive cells were counted by capturing live images (Fig. 2). SG formation was observed as early as 0.5 h after exposure to As^3+^ in CHOG3BP1 cells with a higher number of SG-positive cells at 300 µM As^3+^ compared to 100 µM As^3+^, suggesting that SG formation might contribute to higher viability in cells exposed to 300 µM As^3+^. SGs were not formed clearly either by Sb^3+^, Cd^2+^, Se^4+^, Cu^2+^, Ag^+^, or PAO (Supplementary Fig. 3). Additionally, CHOG3BP1 cells did not form SGs when exposed to 30 µM As^3+^. Pre-treatment of cells with BSO significantly reduced cellular GSH concentrations and increased sensitivity to As^3+^. However, SGs were not formed by 30 µM As^3+^ (Fig. 3), indicating that GSH depletion did not enhance As^3+^-induced SG formation, if any, and suggesting that high concentrations of As^3+^ (>100 µM) are required for SG formation.

**Fig. 2.**
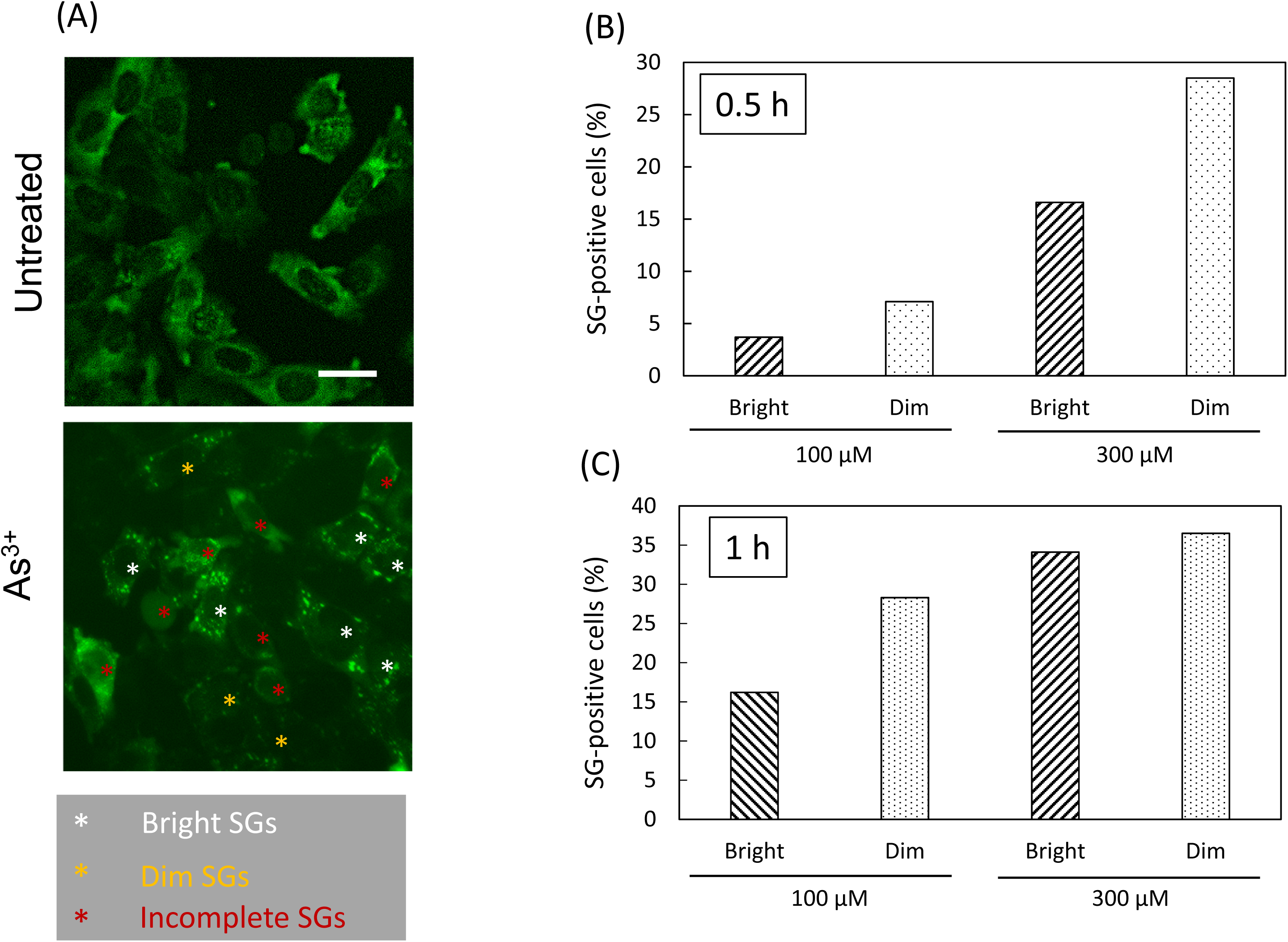
Formation of SGs in As^3+^-exposed CHOG3BP1 cells. Fluorescent photomicrographs of untreated CHOG3BP1 cells and the cells treated with 300 µM As^3+^ for 0.5 h (A). Cells were categorized into three types: Cells with five or more than five bright (white asterisk) and dim (yellow asterisk) SGs without diffuse green fluorescence in the cytoplasm were counted as SG-positive. Cells with diffuse staining or with less than 5 granules were categorized as incomplete (red asterisk). Cells were exposed to 100 or 300 µM As^3+^ for 0.5 (B) or 1 h (C), and cells in each category were counted. Scale bar: 20 μm.

**Fig. 3.**
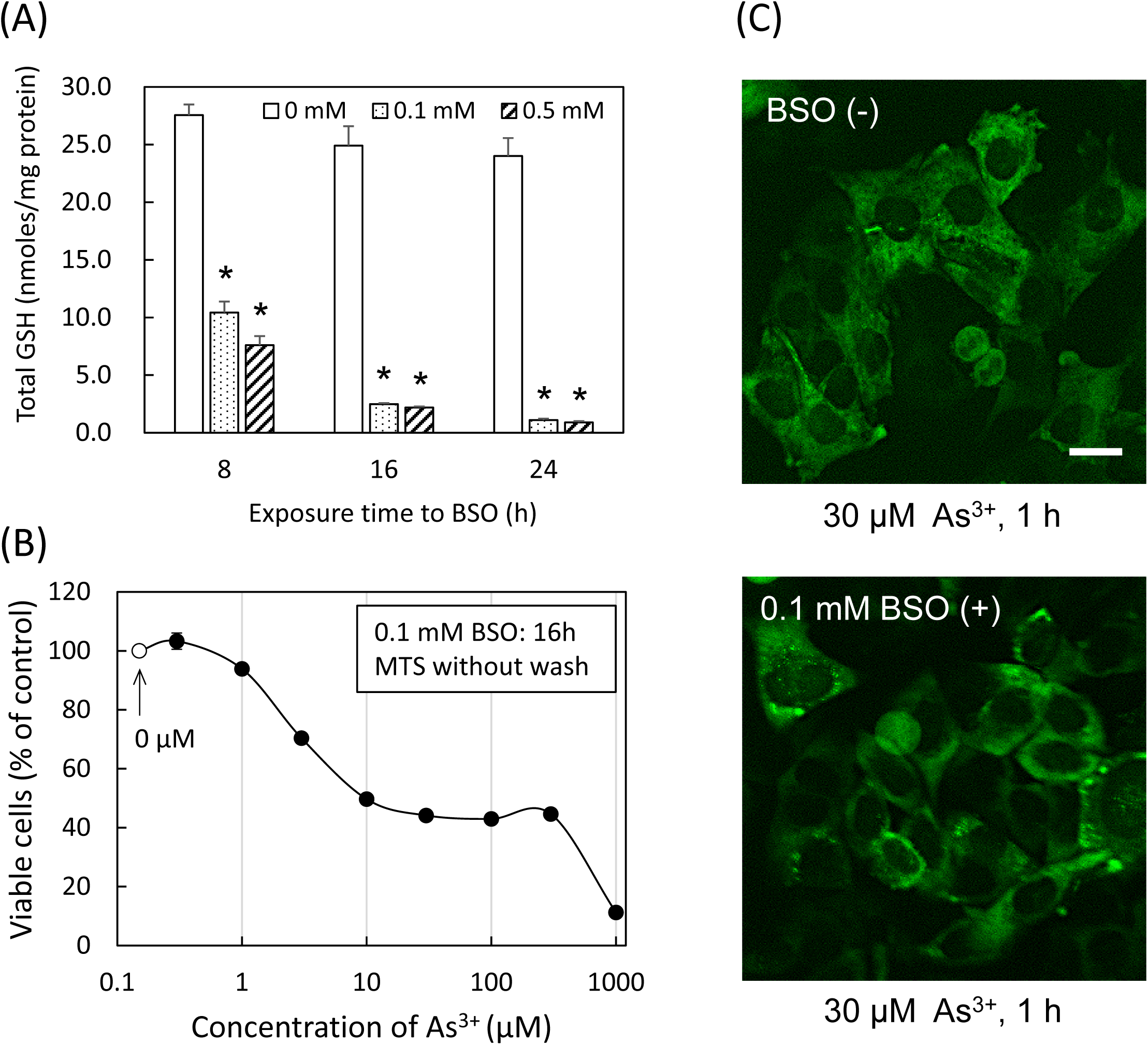
Effects of BSO on GSH concentrations (A), As^3+^-induced cytotoxicity (B), and induction of SGs (C) in CHOG3BP1 cells. (A) CHOG3BP1 cells were exposed to 0.1, and 0.5 mM BSO for 8, 16, and 24 h for measurements of GSH concentrations. (B) Cells pre-treated with 0.1 mM BSO for 16 h were exposed to As^3+^ for 24 h, and viability was assayed by MTS without washing. (C) Cells untreated or pre-treated with 0.1 mM BSO for 16 h were exposed to 30 µM As^3+^ for 1 h. Data are presented as mean ± SEM (N=4). *, Significantly different from untreated (0 mM BSO). Scale bar: 20 μm.

### Cytotoxicity of As^3+^ in leukemia cells and SG formation in JurG3BP1 cells

We further investigated the non-canonical cytotoxic effects of As^3+^ using various cell lines. Both JurG3BP1 (Fig. 4) and Jurkat cells (Supplementary Fig. 4) exhibited similar non-canonical sensitivity to As^3+^ and a concentration-dependent sensitivity to Cd^2+^, akin to CHOG3BP1 and CHO-K1 cells. FBS did not affect the sensitivity of Jurkat cells to As^3+^ (Supplementary Fig. 4). In JurG3BP1 cells, BSO reduced GSH concentration and increased sensitivity to As^3+^. However, the biphasic cytotoxicity feature became less evident in the presence of BSO (Fig. 4). To explore whether the observed pro-survival tendency at 300 µM As^3+^ is a general phenomenon, we examined two other human leukemia cell lines. THP-1 and HL60 cells also demonstrated non-canonical survival responses to As^3+^, although cytotoxicity reversed at 1000 µM As^3+^ in these cell lines (Supplementary Fig. 5), suggesting cell-type dependency for optimal non-canonical concentrations of As^3+^.

**Fig. 4.**
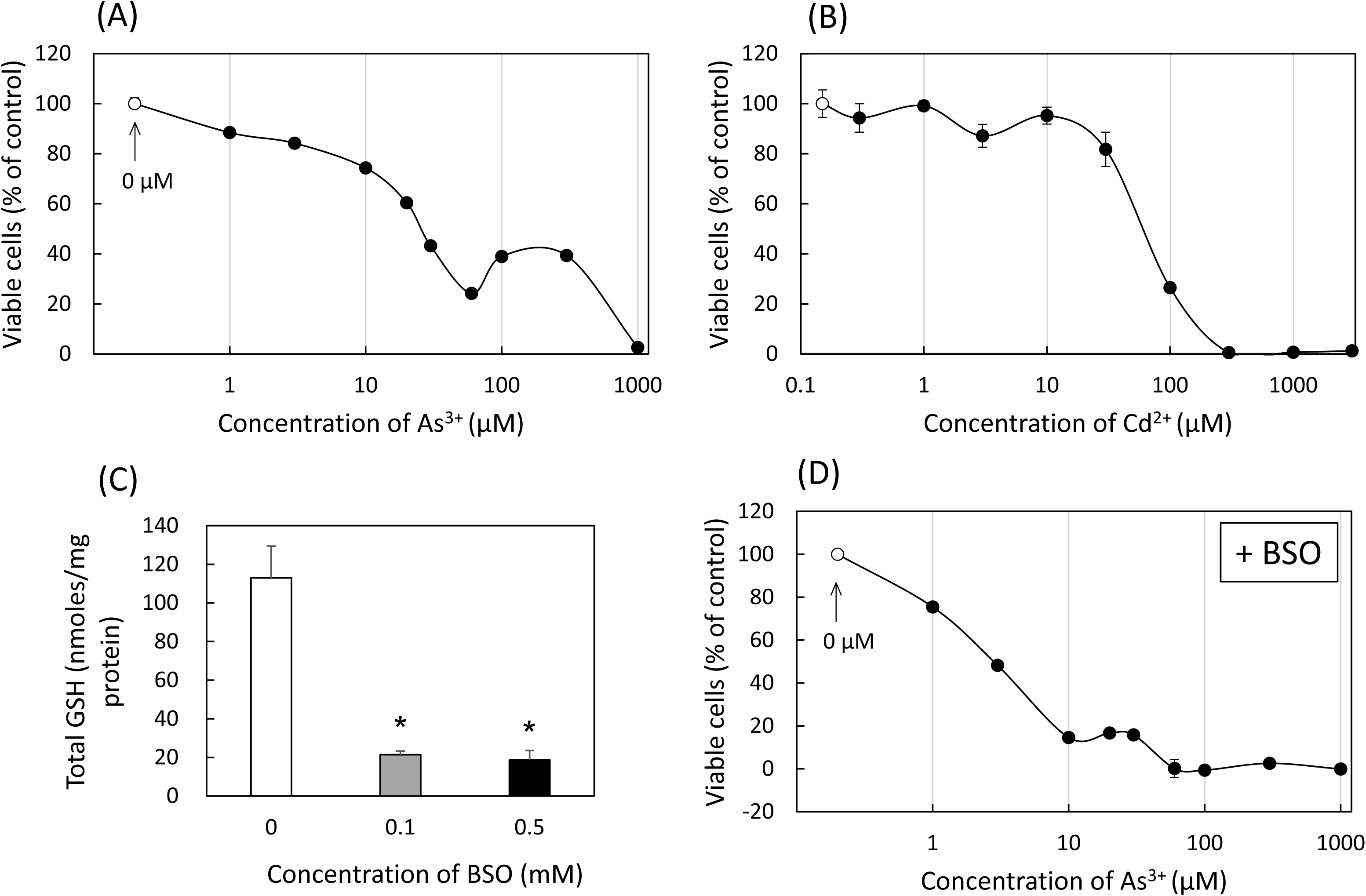
Cytotoxicity of As^3+^ (A) and Cd^2+^ (B) in JurG3BP1 cells, and effects of BSO on GSH concentrations (C) and As^3+^-induced cytotoxicity (D). (A and B) JurG3BP1 cells were exposed to pre-determined concentrations of As^3+^ or Cd^2+^ for 24 h. EC_50_ value for Cd^2+^ was calculated as 67 μM. (C) Cells were exposed to 0.1 or 0.5 mM BSO for 16 h. (D) Cells pre-treated with 0.1 mM BSO for 16 h were exposed to As^3+^ for 24 h. The cell viability was assayed by MTS. Data are presented as mean ± SEM (N=5 for Cd^2+^, otherwise N=4). *, Significantly different from control.

Jurkat cells were exposed to pre-determined concentrations of As^3+^ for 6 h and then cultured further in As^3+^-free culture medium to examine reproducibility of the non-canonical survival response under different exposure conditions. Cells exposed to 300 µM As^3+^ resumed growth faster than those exposed to 100 µM As^3+^ (Fig. 5A). As^3+^ induced cell death partially through programmed pathways, as evidenced by slight but significant reduction in cytotoxicity in Z-VAD (an apoptosis inhibitor) and necrostatin-1 (a necroptosis inhibitor) (Fig. 5B). Heat treatment, hypertonic stress with sorbitol, and As^3+^ exposure induced SGs in JurG3BP1 cells. DTT, a strong reducing agent, induced SGs in JurG3BP1 cells at 1mM (Fig. 6), suggesting that oxidative stress is not requisite for SG formation.

**Fig. 5.**
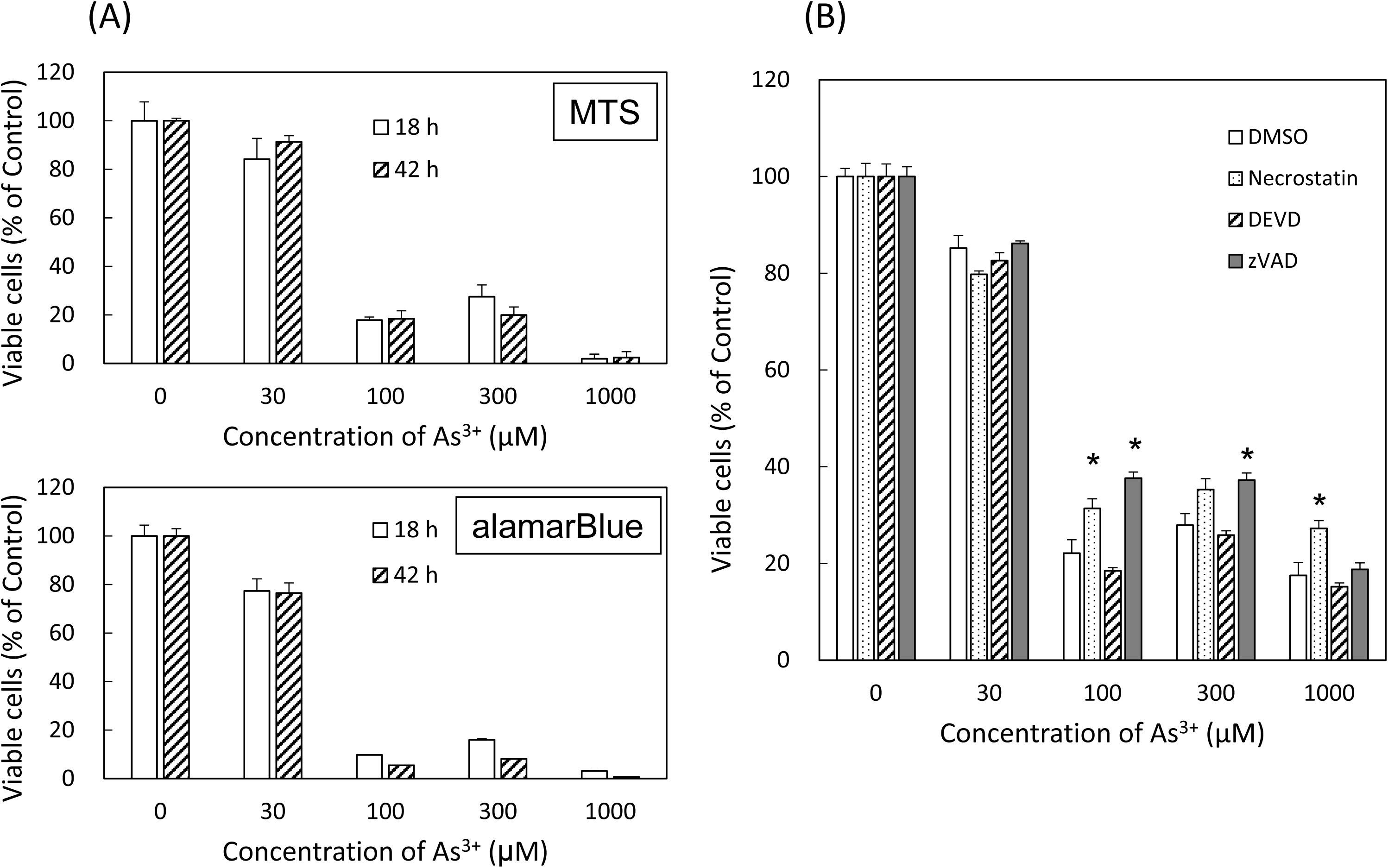
Recovery from cytotoxic effects of As^3+^ (A) and effects of cell death inhibitors on As^3+^-induced cytotoxicity in Jurkat cells (B). (A) Jurkat cells were exposed to increasing concentrations of As^3+^ for 6 h were washed and further cultured in fresh complete culture medium for 18 and 42 h. Cell viability was assayed by both MTS and alamarBlue methods. (B) Cells were exposed to As^3+^ for 24 h with or without inhibitors (10 µM necrostatin-1, 10 µM Ac-DEVD-CHO, 20 µM Z-VAD-FMK, or 0.1% DMSO). The inhibitors were added 1 h before addition of As^3+^. Viability was assayed by MTS. As^3+^-induced cytotoxicity was reduced by Z-VAD-FMK (a pan-caspase inhibitor), but not by Ac-DEVD-CHO (a caspase-3 inhibitor). Data are presented as mean ± SEM (N=4). *, Significantly different from control.

**Fig. 6.**
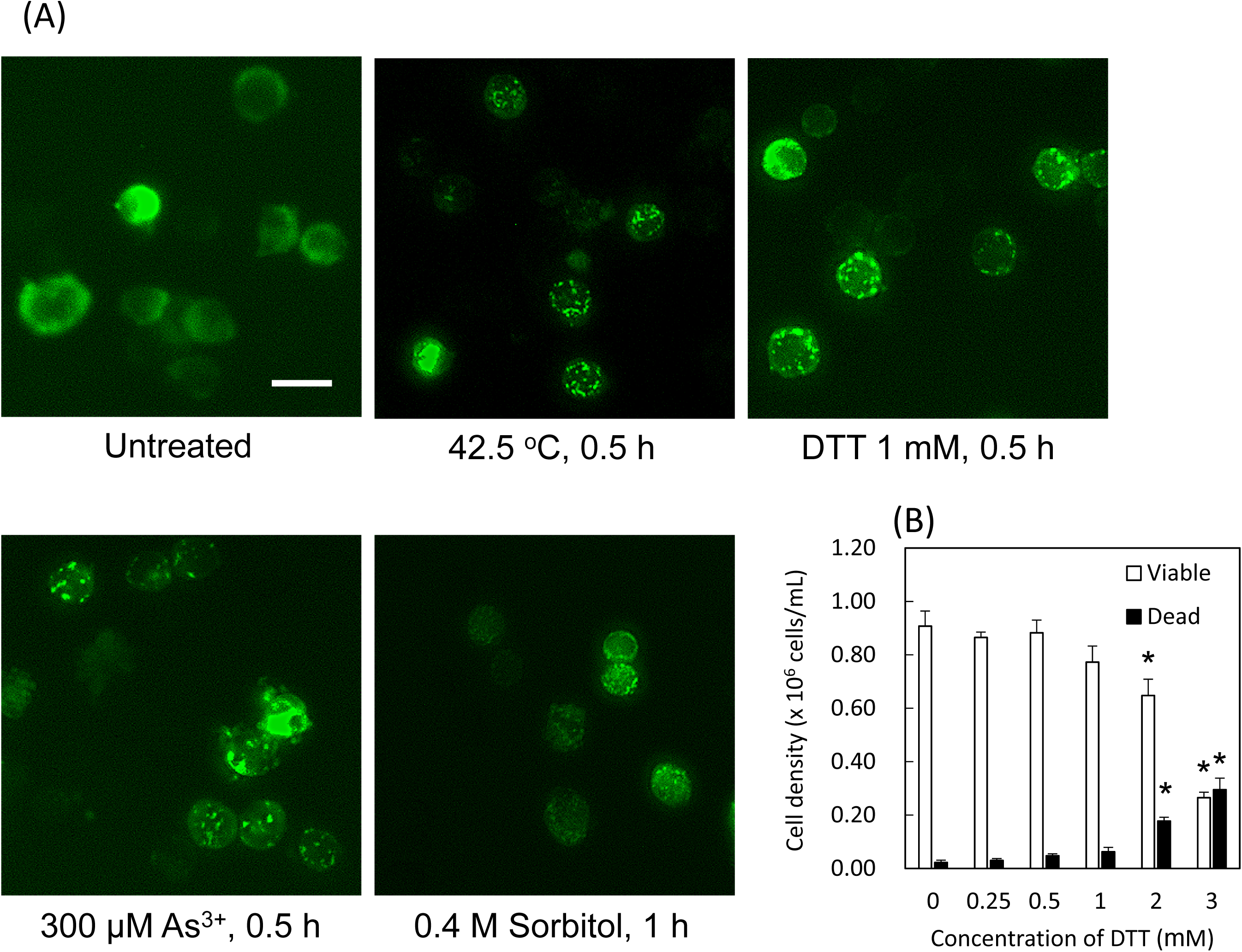
Induction of SGs in JurG3BP1 cells under various stress conditions (A), and effects of DTT on Jurkat cell viability (B). (A) JurG3BP1 cells were exposed to 42.5 ℃ heat shock, 300 µM As^3+^, 0.4 M sorbitol, or 1 mM DTT for specified times. (B) Jurkat cells were exposed to pre-determined concentrations of DTT for 24 h for measurement of cell viability. Viability was assayed by trypan blue dye exclusion because colorimetric methods were not applicable for the strong reducing property of DTT. Data are presented as mean ± SEM (N=4). *, Significantly different from the untreated group (0 mM DTT). Scale bar: 20 μm.

### Phosphorylation of eIF2α in response to As^3+^ and other stimuli

Fig. 7 shows phosphorylation of eIF2α in response to As^3+^ compared with Cd^2+^, hypertonic, and heat shock. In CHOG3BP1 cells, As^3+^ increased eIF2α phosphorylation levels which were comparable to Cd^2+^ but lower than those induced by hypertonic shock (Fig. 7A). In JurG3BP1 cells, both 300 µM As^3+^ and heat shock similarly elevated eIF2α phosphorylation levels. Exposure to 100 µM As^3+^ and 100 µM Cd^2+^ resulted in nearly the same phosphorylation level (Fig. 7B).

**Fig. 7.**
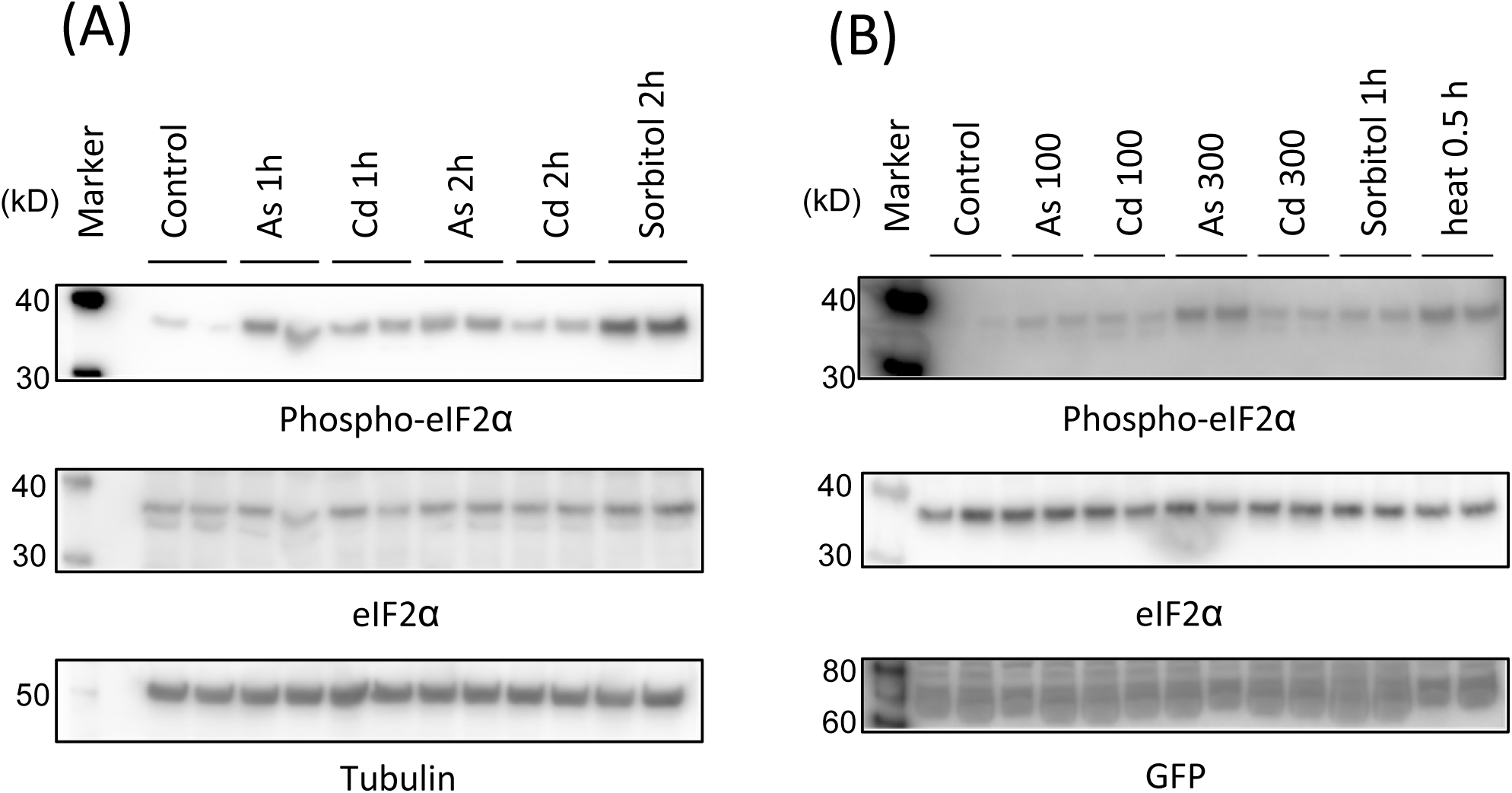
Western blot analyses for eIF2α phosphorylation in CHOG3BP1 (A) and JurG3BP1 cells (B). (A) CHOG3BP1 cells were exposed to 300 µM As^3+^ or 300 µM Cd^2+^ for 1 or 2 h, or 0.4 M sorbitol for 2 h. (B) JurG3BP1 cells were exposed 100 or 300 µM As^3+^ or Cd^2+^ for 1 h, 0.4 M sorbitol for 1 h, or heat shock (42.5 ℃) for 0.5 h. After exposure cells were lyzed with cold RIPA solution, and the supernatant of the lysate was analyzed by western blotting. Lysates from two separate wells were analyzed.

### SUMO proteins did not contribute to SG formation

Endogenous SUMO molecules have not been consistently detected in SGs thus far, despite indication that SUMOylation of SG-associated proteins may influence their recruitment to SGs [20]. We investigated whether overexpressed SUMO molecules contribute to SG formation through covalent (SUMOylation) or non-covalent (SUMO-SIM interaction) binding, or both. mCherry-tagged wild-type SUMO2 did not co-localize with SGs even under As^3+^, hypertonic, and heat shock conditions (Fig. 8A). Notably, mCherry-tagged wild-type SUMO2 formed granular structures in the nuclei under heat shock, whereas SGs form in the cytoplasm. To explore the potential non-covalent interaction of SUMO molecules with SG proteins, we employed SUMOylation-disabled mutant SUMO2 and a SUMO E1 inhibitor, ML792. The ΔGG mutant SUMO2, which diffusely distribute throughout the cells, did not co-localize with SGs following stimulation with As^3+^, hypertonicity, or heat shock (Fig. 8B). ML792 treatment resulted in cytoplasmic release of SUMO2 but did not lead to its involvement in SGs (Supplementary Fig. 6).

**Fig. 8.**
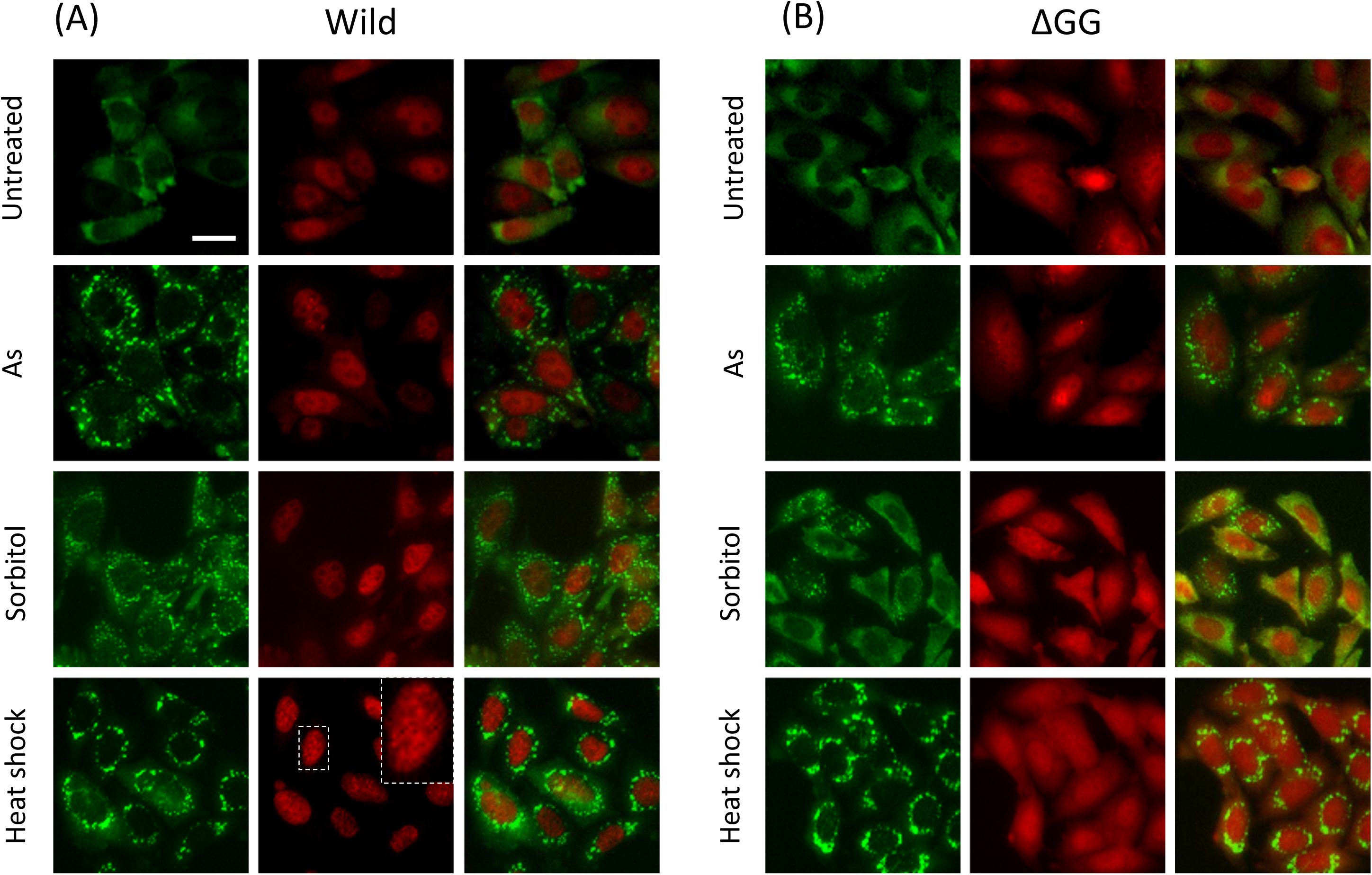
SUMO2 involvement in SG formation. CHOG3BP1 cells expressing wild-type (A) and C-terminal diglycine-deleted (ΔGG) mCherry-tagged SUMO2 (B) were treated with 300 µM As^3+^ for 30 min, 0.4 M sorbitol for 2 h, or heat shock for 45 min. The enlarged image is shown on the right upper corner.

### Changes in promyelocytic leukemia proteins (PML), SUMO, and ubiquitin in response to As^3+^ and hypertonic stress

As^3+^ rapidly reduces the solubility of PML proteins and enhances their SUMOylation at a concentration of 3 µM [8, 9, 21, 22]. We investigated intracellular dynamics of SUMO and PML proteins after exposure to As^3+^ and 0.4 M sorbitol to explore whether SG formation alters the biochemical properties of PML proteins. HEKG3BP1 cells were utilized instead of CHOG3BP1 cells due to the availability of an anti-human PML antibody for the detection of endogenous PML in HEK cells. Exposure to 50 µM As^3+^ and 0.4 M sorbitol formed bright large and small SGs, respectively. Cells exposed to 3 µM As^3+^, 0.4 M sorbitol, or both were lyzed with ice-cold RIPA buffer for western blot analysis. Exposure to 3 µM As^3+^ induced solubility shift and SUMOylation of PML proteins. Hypertonic shock with 0.4 M sorbitol increased levels of SUMOylated and ubiquitinated proteins and partially altered their solubility. However, sorbitol did not affect the solubility or SUMOylation level of PML proteins, suggesting different signaling pathways between SG formation and responses of PML proteins in As^3+^-exposed cells (Fig. 9).

**Fig. 9.**
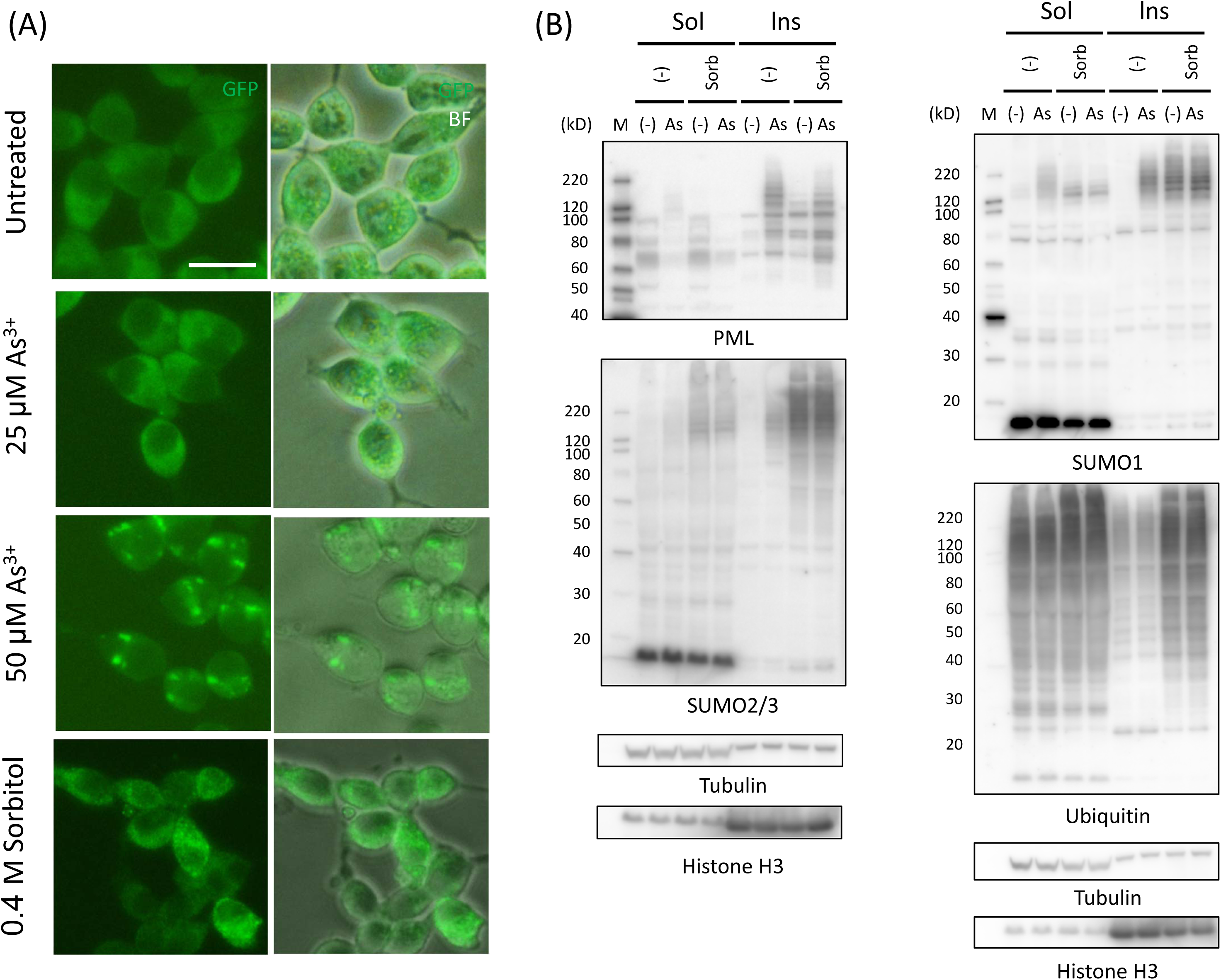
SG formation in HEKG3BP1 cells after exposure to As^3+^ or sorbitol (A), and western blot analyses for PML, SUMO, and ubiquitin proteins (B). (A) HEKG3BP1 cells were exposed to 25 and 50 μM As^3+^ or 0.4 M sorbitol for 1 h. Scale bar: 20 μm. (B) Cells were lysed in ice-cold RIPA solution and the lysates were separated into RIPA-soluble and RIPA-insoluble fractions. After blotting, the membranes were probed for PML, SUMO2/3, SUMO1 and ubiquitin. The membranes were finally probed with premixed HRP-tagged anti-tubulin and HRP-tagged anti histone H3 antibodies.

## Discussion

### Cytotoxic effects of As^3+^

The detachment of adherent cells from culture plates is considered to be an indicative of cell death [23]. Therefore, washing wells of assay plates is standard practice for the colorimetric cytotoxicity assay using adherent cells. Cells with rounding up or less adherent morphology are removed by vigorous washing of cell monolayers even though these cells are alive and functional after removal of stress. In the present study, we observed that CHO and Jurkat cells demonstrated greater resilience to As^3+^-intoxication at 300 µM compared to 100 µM. Other cell lines such as THP-1 and HL60 showed resistance to As^3+^-induced cytotoxic effects at 1000 µM. Notably, this non-canonical concentration-cytotoxicity relationship appears specific to As^3+^, as other toxic metals killed cells in a typical concentration-dependent manner. To our best knowledge, only one previous study has reported a similar non-canonical or biphasic concentration-survival curve in As^3+^-induced cytotoxicity assays specifically noting a peak at 3 mM in HeLa cells [15]. The uptake of As^3+^ occurs through aquaglyceroporins [24], followed by conjugation with GSH [25] and subsequent methylation [26]. The excretion of arsenic and its metabolites via multidrug resistance-associated proteins [27, 28] influences the intracellular concentrations of reactive arsenic species [29], which likely contribute to the observed differences in optimal As^3+^ concentrations for resilience across various cell types.

Type I stresses such as hypoxia, heat shock, and As^3+^ typically induce SG formation. RACK1, sequestered within SGs, suppresses MTK1-SAPK activation pathway, and mitigates apoptosis under condition of Type I stress. Conversely, Type II stressors like irradiation and genotoxic drugs activate p38 and JNK MAPK pathways, leading cells towards apoptosis [30]. The complexity of cell death mechanism in As^3+^-exposed cells is evident from our findings that the pan-caspase inhibitor Z-VAD and a necroptosis inhibitor necrostatin-1 only partially reduced As^3+^-induced cytotoxicity (Fig. 5). This suggests that Type I stress responses may interfere with apoptosis pathways and contribute to the observed resilience.

### Triggers of SG formation

Exposure to 30-1000 µM As^3+^ for 30-45 min is commonly used to induce SGs. It has been reported that exposure to As^3+^ leads to inhibition of translation initiation and disappearance of polysomes [16]. As^3+^ is known to react with cysteine residues and induce oxidative stress [9], yet oxidation of intracellular sulfhydryl groups alone does not always lead to SG formation, as demonstrated by the induction of SGs by DTT, a thiol-base antioxidant (Fig. 6). The exact mechanism of DTT-induced SG formation remains unclear [31], but it is plausible that As^3+^ and DTT induce SGs via reacting with cysteine residues of proteins [32, 33], reducing disulfide, and promoting the assembly of unfolded, aggregation-prone proteins into SGs.

Phosphorylation of eIF2α hampers formation of the eIF2-GTP-tRNA^Met^ ternary complex, thereby stalling translation initiation, a prerequisite for SG formation. [16]. Interestingly, microinjection of mRNA and ssDNA, but not dsDNA induced SGs, highlighting the requirement for active mRNA translocation in SG assembly. Our findings also revealed that Cd^2+^ increases phosphorylation of eIF2α, yet does not lead to SG formation, indicating that eIF2α phosphorylation alone is not sufficient to induce SGs.

### Modulation of cytotoxicity in As^3+^-exposed cells, and specific cellular events induced by As^3+^

SG formation represents a pro-survival response in cells exposed to severe stresses such as heat, hypertonicity, and As^3+^. The present study unequivocally demonstrated that survival curves of As^3+^-exposed cells exhibited a non-canonical, concentration-independent pattern at concentrations where SGs were induced by As^3+^. As^3+^ is stabilized by conjugation by GSH, forming arsenic triglutathione [34]. Consequently, depletion of intracellular GSH renders cells more susceptible to As^3+^, underscoring a protective role of GSH against oxidative stress. Figs. 3 and 4 show that BSO reduced intracellular GSH levels and increased susceptibility to As^3+^, diminishing the non-canonical survival curve in both CHOG3BP1 and JurG3BP1 cells. Notably, As^3+^-induced SG formation was not enhanced by BSO treatment, suggesting that GSH may not directly influence SG formation.

Two distinct cellular events appear to differentiate As^3+^ from other toxic metal or metalloid ions; the induction of SG formation in the cytoplasm and biochemical alterations of PML in the nucleoplasm. Specifically, SGs formation is specific to As^3+^ among toxic metals, while solubility loss and SUMOylation of PML proteins are unique to As^3+^ and Sb^3+^ exposure [8, 10]. We investigated whether SG formation and biochemical changes of PML corelate in As^3+^-exposed cells. Hypertonic stress with 0.4 M sorbitol increased SUMOylation and ubiquitination levels of proteins, partially converting them into cold-RIPA insoluble forms (Fig 9). However, PML proteins were unaffected, if any, by sorbitol. In contrast, exposure to 3 µM As^3+^ selectively induced solubility loss and SUMOylation of PML proteins, indicating a specific response of PML to As^3+^ that was not observed with 0.4 M sorbitol despite both stressors’ ability to induce SGs.

It is widely accepted that environmental stresses such as heat shock, hypertonicity, and hydrogen peroxide induce SUMOylation in mammalian cells [35] and the SUMOylation of poly (ADP-ribose) polymerase (PARP-1) is implicated in apoptosis [36]. Most wild-type SUMO2 molecules were present in the nuclei, where heat shock transformed SUMO2 into dot-like structures (Fig. 8A). These changes were attributed to increased SUMOylation levels, as demonstrated by the lack of this response in ΔGG mutant SUMO2 expressing cells (Fig. 8B). SUMO molecules may be involved in phase separation with PML [37] or preventing the accumulation of unfolded proteins into insoluble aggregates [38]. Notably, SG formation occurred independently of functional (wild-type) and unfunctional (ΔGG) SUMO2 expressions (Fig. 8), indicating that SUMO molecules do not directly influence SG formation despite their overexpression.

## Conclusions

Arsenite (As^3+^) induced less cytotoxicity at 300 µM compared to 100 µM As^3+^ in CHO and Jurkat cells. Rapid SG formation at 300 µM As^3+^ may function as a pro-survival mechanism, contributing the observed non-canonical survival curves. The present study also warrants that viable cells are inadvertently discarded if cell monolayers are washed vigorously in cytotoxicity assays.

## Supporting Information

A source table and uncropped western images.

## Supporting information

Uncropped western image (Fig.7)

Uncropped western image (Fig.9)

## Acknowledgments

This work was partially supported by JSPS (#18H03043).

## References

1. IPCS. Arsenic: World Health Organization; 1981.

2. IPCS. Arsenic and arsenic compounds. Geneva: World Health Organization; 2001.

3. Fox E, Razzouk BI, Widemann BC, Xiao S, O’Brien M, Goodspeed W, et al. Phase 1 trial and pharmacokinetic study of arsenic trioxide in children and adolescents with refractory or relapsed acute leukemia, including acute promyelocytic leukemia or lymphoma. Blood. 2008;111(2):566–73. doi: 10.1182/blood-2007-08-107839. PubMed PMID: 17959855; PubMed Central PMCID: PMC2200837.

4. Berenson J, Matous J, Swift R, Mapes R, Morrison B, Yeh H. A phase I/II study of arsenic trioxide/bortezomib/ascorbic acid combination therapy for the treatment of relapsed or refractory multiple myeloma. Clin Cancer Res. 2007;13(6):1762–8. doi: 10.1158/1078-0432.CCR-06-1812. PubMed PMID: 17363530.

5. Hirano S. Biotransformation of arsenic and toxicological implication of arsenic metabolites. Arch Toxicol. 2020;94(8):2587–601. doi: 10.1007/s00204-020-02772-9. PubMed PMID: 32435915.

6. Sato M, Kondoh M. Recent studies on metallothionein: protection against toxicity of heavy metals and oxygen free radicals. Tohoku J Exp Med. 2002;196(1):9–22. doi: 10.1620/tjem.196.9. PubMed PMID: 12498322.

7. Suzuki KT, Takahara T, Watanabe H, Nishikawa M, Yamamura M, Murakami M. Effect of pretreatment with cadmium/cysteine or metallothionein on accumulation of cadmium challenged with either complexes. Arch Toxicol. 1986;58(4):261–4. doi: 10.1007/BF00297117. PubMed PMID: 3718230.

8. Hirano S, Tadano M, Kobayashi Y, Udagawa O, Kato A. Solubility shift and SUMOylaltion of promyelocytic leukemia (PML) protein in response to arsenic(III) and fate of the SUMOylated PML. Toxicol Appl Pharmacol. 2015;287(3):191–201. doi: 10.1016/j.taap.2015.05.018. PubMed PMID: 26049103.

9. Hirano S, Watanabe T, Kobayashi Y. Effects of arsenic on modification of promyelocytic leukemia (PML): PML responds to low levels of arsenite. Toxicol Appl Pharmacol. 2013;273(3):590–9. doi: 10.1016/j.taap.2013.10.004. PubMed PMID: 24135626.

10. Muller S, Miller WH, Jr., Dejean A. Trivalent antimonials induce degradation of the PML-RAR oncoprotein and reorganization of the promyelocytic leukemia nuclear bodies in acute promyelocytic leukemia NB4 cells. Blood. 1998;92(11):4308–16. PubMed PMID: 9834237.

11. Fujimura K, Sasaki A, Anderson P. Selenite targets eIF4E-binding protein-1 to inhibit translation initiation and induce the assembly of non-canonical stress granules. Nucleic Acids Res. 2012;40(16):8099–110. doi: 10.1093/nar/gks566. PubMed PMID: 22718973; PubMed Central PMCID: PMC3439927.

12. Aulas A, Lyons S, Fay M, Anderson P, Ivanov P. Nitric oxide triggers the assembly of “type II” stress granules linked to decreased cell viability. Cell Death Dis. 2018;9(11):1129. doi: 10.1038/s41419-018-1173-x. PubMed PMID: 30425239; PubMed Central PMCID: PMC6234215.

13. Hofmann S, Kedersha N, Anderson P, Ivanov P. Molecular mechanisms of stress granule assembly and disassembly. Biochim Biophys Acta Mol Cell Res. 2020;1868(1):118876. doi: 10.1016/j.bbamcr.2020.118876. PubMed PMID: 33007331.

14. Riggs C, Kedersha N, Ivanov P, Anderson P. Mammalian stress granules and P bodies at a glance. J Cell Sci. 2020;133(16). doi: 10.1242/jcs.242487. PubMed PMID: 32873715.

15. Glass L, Wente SR. Gle1 mediates stress granule-dependent survival during chemotoxic stress. Adv Biol Regul. 2019;71:156–71. doi: 10.1016/j.jbior.2018.09.007. PubMed PMID: 30262214; PubMed Central PMCID: PMC6347492.

16. Bounedjah O, Desforges B, Wu T, Pioche-Durieu C, Marco S, Hamon L, et al. Free mRNA in excess upon polysome dissociation is a scaffold for protein multimerization to form stress granules. Nucleic Acids Res. 2014;42(13):8678–91. doi: 10.1093/nar/gku582. PubMed PMID: 25013173; PubMed Central PMCID: PMC4117795.

17. Bussi C, Mangiarotti A, Vanhille-Campos C, Aylan B, Pellegrino E, Athanasiadi N, et al. Stress granules plug and stabilize damaged endolysosomal membranes. Nature. 2023;623(7989):1062–9. doi: 10.1038/s41586-023-06726-w. PubMed PMID: 37968398; PubMed Central PMCID: PMC10686833.

18. Samir P, Kesavardhana S, Patmore DM, Gingras S, Malireddi RKS, Karki R, et al. DDX3X acts as a live-or-die checkpoint in stressed cells by regulating NLRP3 inflammasome. Nature. 2019;573(7775):590–4. doi: 10.1038/s41586-019-1551-2. PubMed PMID: 31511697; PubMed Central PMCID: PMC6980284.

19. Takahashi M, Higuchi M, Matsuki H, Yoshita M, Ohsawa T, Oie M, et al. Stress granules inhibit apoptosis by reducing reactive oxygen species production. Mol Cell Biol. 2013;33(4):815–29. doi: 10.1128/MCB.00763-12. PubMed PMID: 23230274; PubMed Central PMCID: PMC3571346.

20. Keiten-Schmitz J, Roder L, Hornstein E, Muller-McNicoll M, Muller S. SUMO: Glue or Solvent for Phase-Separated Ribonucleoprotein Complexes and Molecular Condensates? Front Mol Biosci. 2021;8:673038. doi: 10.3389/fmolb.2021.673038. PubMed PMID: 34026847; PubMed Central PMCID: PMC8138125.

21. Hirano S, Udagawa O. SUMOylation regulates the number and size of promyelocytic leukemia-nuclear bodies (PML-NBs) and arsenic perturbs SUMO dynamics on PML by insolubilizing PML in THP-1 cells. Arch Toxicol. 2022;96(2):545–58. doi: 10.1007/s00204-021-03195-w. PubMed PMID: 35001170.

22. Hirano S, Udagawa O, Kobayashi Y, Kato A. Solubility changes of promyelocytic leukemia (PML) and SUMO monomers and dynamics of PML nuclear body proteins in arsenite-treated cells. Toxicol Appl Pharmacol. 2018;360:150–9. doi: 10.1016/j.taap.2018.10.001. PubMed PMID: 30292834.

23. Hofmann S, Cherkasova V, Bankhead P, Bukau B, Stoecklin G. Translation suppression promotes stress granule formation and cell survival in response to cold shock. Mol Biol Cell. 2012;23(19):3786–800. doi: 10.1091/mbc.E12-04-0296. PubMed PMID: 22875991; PubMed Central PMCID: PMC3459856.

24. Liu Z, Shen J, Carbrey JM, Mukhopadhyay R, Agre P, Rosen BP. Arsenite transport by mammalian aquaglyceroporins AQP7 and AQP9. Proc Natl Acad Sci U S A. 2002;99(9):6053–8. Epub 2002 Apr 23. PubMed PMID: 11972053.

25. Kobayashi Y, Hirano S. Effects of endogenous hydrogen peroxide and glutathione on the stability of arsenic metabolites in rat bile. Toxicol Appl Pharmacol. 2008;232(1):33–40. doi: 10.1016/j.taap.2008.06.003. PubMed PMID: 18619986.

26. Hayakawa T, Kobayashi Y, Cui X, Hirano S. A new metabolic pathway of arsenite: arsenic-glutathione complexes are substrates for human arsenic methyltransferase Cyt19. Arch Toxicol. 2005;79(4):183–91. doi: 10.1007/s00204-004-0620-x. PubMed PMID: 15526190.

27. Dietrich CG, Ottenhoff R, de Waart DR, Elferink R. Role of MRP2 and GSH in intrahepatic cycling of toxins. Toxicology. 2001;167(1):73–81. PubMed PMID: ISI:000171355800007.

28. Kala SV, Neely MW, Kala G, Prater CI, Atwood DW, Rice JS, et al. The MRP2/cMOAT transporter and arsenic-glutathione complex formation are required for biliary excretion of arsenic. J Biol Chem. 2000;275(43):33404–8.

29. Suzuki KT, Tomita T, Ogra Y, Ohmichi M. Glutathione-conjugated arsenics in the potential hepato-enteric circulation in rats. Chemical Research in Toxicology. 2001;14(12):1604–11. PubMed PMID: ISI:000172934700006.

30. Arimoto K, Fukuda H, Imajoh-Ohmi S, Saito H, Takekawa M. Formation of stress granules inhibits apoptosis by suppressing stress-responsive MAPK pathways. Nat Cell Biol. 2008;10(11):1324–32. doi: 10.1038/ncb1791. PubMed PMID: 18836437.

31. Basu M, Courtney S, Brinton M. Arsenite-induced stress granule formation is inhibited by elevated levels of reduced glutathione in West Nile virus-infected cells. PLoS Pathog. 2017;13(2):e1006240. doi: 10.1371/journal.ppat.1006240. PubMed PMID: 28241074; PubMed Central PMCID: PMC5344523.

32. Kitchin KT, Wallace K. Arsenite binding to synthetic peptides based on the Zn finger region and the estrogen binding region of the human estrogen receptor-alpha. Toxicol Appl Pharmacol. 2005;206(1):66–72. doi: 10.1016/j.taap.2004.12.010. PubMed PMID: 15963345.

33. Mizumura A, Watanabe T, Kobayashi Y, Hirano S. Identification of arsenite-and arsenic diglutathione-binding proteins in human hepatocarcinoma cells. Toxicol Appl Pharmacol. 2010;242(2):119–25. doi: 10.1016/j.taap.2009.10.013. PubMed PMID: 19874836.

34. Watanabe T, Hirano S. Metabolism of arsenic and its toxicological relevance. Arch Toxicol. 2013;87(6):969–79. doi: 10.1007/s00204-012-0904-5. PubMed PMID: 22811022.

35. Saitoh H, Hinchey J. Functional heterogeneity of small ubiquitin-related protein modifiers SUMO-1 versus SUMO-2/3. J Biol Chem. 2000;275(9):6252–8. PubMed PMID: 10692421.

36. Martin N, Schwamborn K, Schreiber V, Werner A, Guillier C, Zhang XD, et al. PARP-1 transcriptional activity is regulated by sumoylation upon heat shock. EMBO J. 2009;28(22):3534–48. doi: 10.1038/emboj.2009.279. PubMed PMID: 19779455; PubMed Central PMCID: PMC2782092.

37. Banani S, Rice A, Peeples W, Lin Y, Jain S, Parker R, et al. Compositional Control of Phase-Separated Cellular Bodies. Cell. 2016;166(3):651–63. doi: 10.1016/j.cell.2016.06.010. PubMed PMID: 27374333; PubMed Central PMCID: PMC4967043.

38. Keiten-Schmitz J, Wagner K, Piller T, Kaulich M, Alberti S, Muller S. The Nuclear SUMO-Targeted Ubiquitin Quality Control Network Regulates the Dynamics of Cytoplasmic Stress Granules. Mol Cell. 2020;79(1):54–67 e7. doi: 10.1016/j.molcel.2020.05.017. PubMed PMID: 32521226.

