## Supplementary Figure for "Formation of stress granules and non-canonical survival responses in arsenite-exposed cells"

### Supplementary Figure Captions

**Supplementary Fig. 1. Photographs for cytotoxicity assay using MTS reagent.** CHOG3BP1 (A) and CHO-K1 cells (B) were exposed to pre-determined concentrations of $As^{3+}$ for 24 h. See also the legend to Fig. 1. Cell monolayers were washed twice with PBS (upper 4 wells) or unwashed (lower 4 wells) before the assay. (C) JurG3BP1 cells pre-treated (lower 4 wells) or untreated (upper 4 wells) with 0.1 mM BSO for 16 h were exposed to $As^{3+}$ for 24 h.

**Supplementary Fig. 2. Cytotoxicity of $Cd^{2+}$ (A), $Cu^{2+}$ (B), $Ag^{+}$ (C), and $Se^{4+}$ (D) in CHOG3BP1 cells.** Cells were exposed to pre-determined concentrations of metal or metalloid salts for 24 h. Cell viability was assayed by MTS without washing. $EC_{50}$ values for $Cd^{2+}$, $Cu^{2+}$, $Ag^{+}$, and $Se^{4+}$ were 18, 118, 33, and 28 μM, respectively. Data are presented as means ± SEM (N=4).

**Supplementary Fig. 3. Fluorescence micrographs of CHOG3BP1 cells exposed to various stressors.** GFP and GFP-bright field overlaid images. Cells were cultured in a glass-bottom dish and pre-cultured overnight in complete F12 culture medium. The cells were left untreated (A) or exposed to 0.5 mM $Sb^{3+}$ for 1 h (B), 125 μM $Cd^{2+}$ for 1 h (C), 1 mM $Se^{4+}$ for 1 h (D), 1 mM $Cu^{2+}$ for 1 h (E), 100 μM $Ag^{+}$ for 0.5 h (F), and 30 μM PAO for 0.5 h (G). Scale bar: 20 μm.

**Supplementary Fig. 4. Cytotoxicity of $As^{3+}$ (A and B) and $Cd^{2+}$ (C) in Jurkat cells.** Cytotoxicity of $As^{3+}$ was assayed in complete RPMI1640 culture medium containing 10% FBS (A) or FBS-free culture medium (B). Cell viability was assayed by almarBlue. $EC_{50}$ for $Cd^{2+}$

was calculated as 71 μM. Note that 10% FBS did not affect the cytotoxic effects of $As^{3+}$. Data are presented as mean ± SEM (N=5 for $Cd^{2+}$, otherwise N=4).

**Supplementary Fig. 5. Cytotoxicity of $As^{3+}$ in THP-1 (A) and HL60 cells (B).** Cells were exposed to pre-determined concentrations of $As^{3+}$ for 24 h in RPMI1640 complete culture medium containing 10% FBS. The cell viability was assayed by alamrBlue. Note that viability was almost completely lost at around 100 μM and gradually increased again peaking at 1000 μM. Data are presented as mean ± SEM (N=4).

**Supplementary Fig. 6. Effects of ML792 on SG formation in CHOG3BP1.** CHOG3BP1 cells expressing wild-type mCherry-tagged SUMO2 were pre-treated with 20 μM ML792 (a SUMO E1 inhibitor) for 4h and then exposed with 300 μM $As^{3+}$ for 1 h. Scale bar: 20 μm.

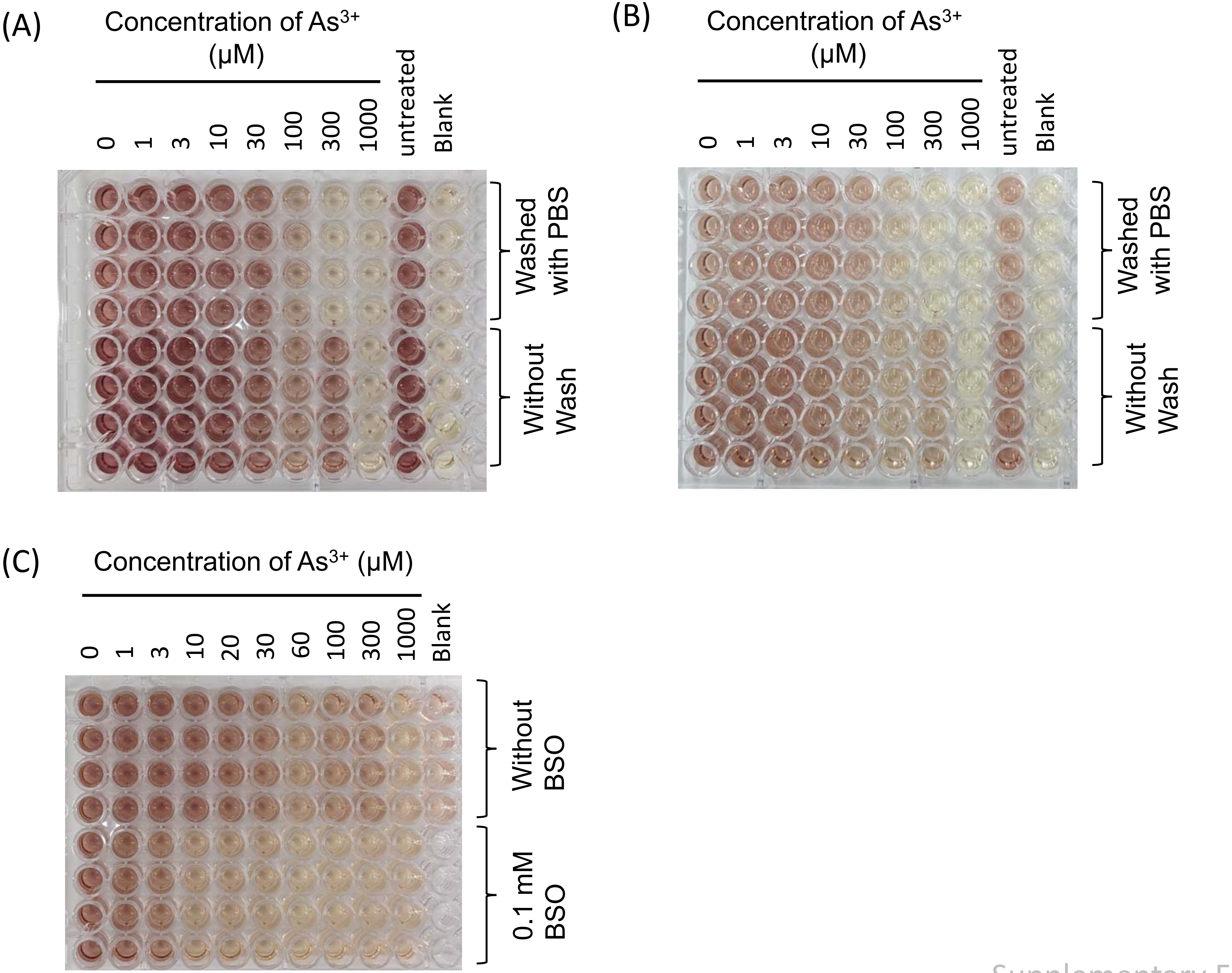


Supplementary Fig. 1

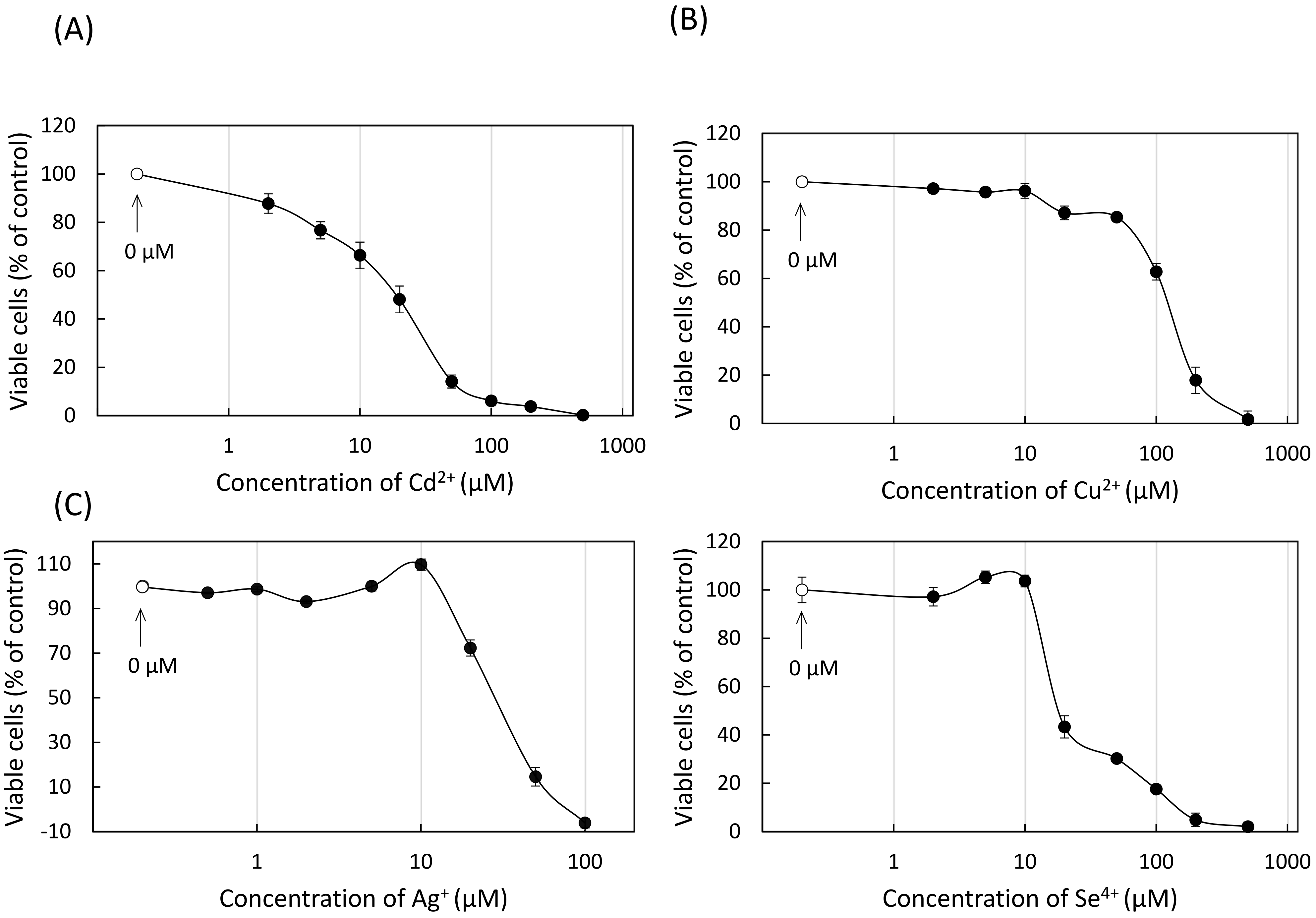


Supplementary Fig. 2

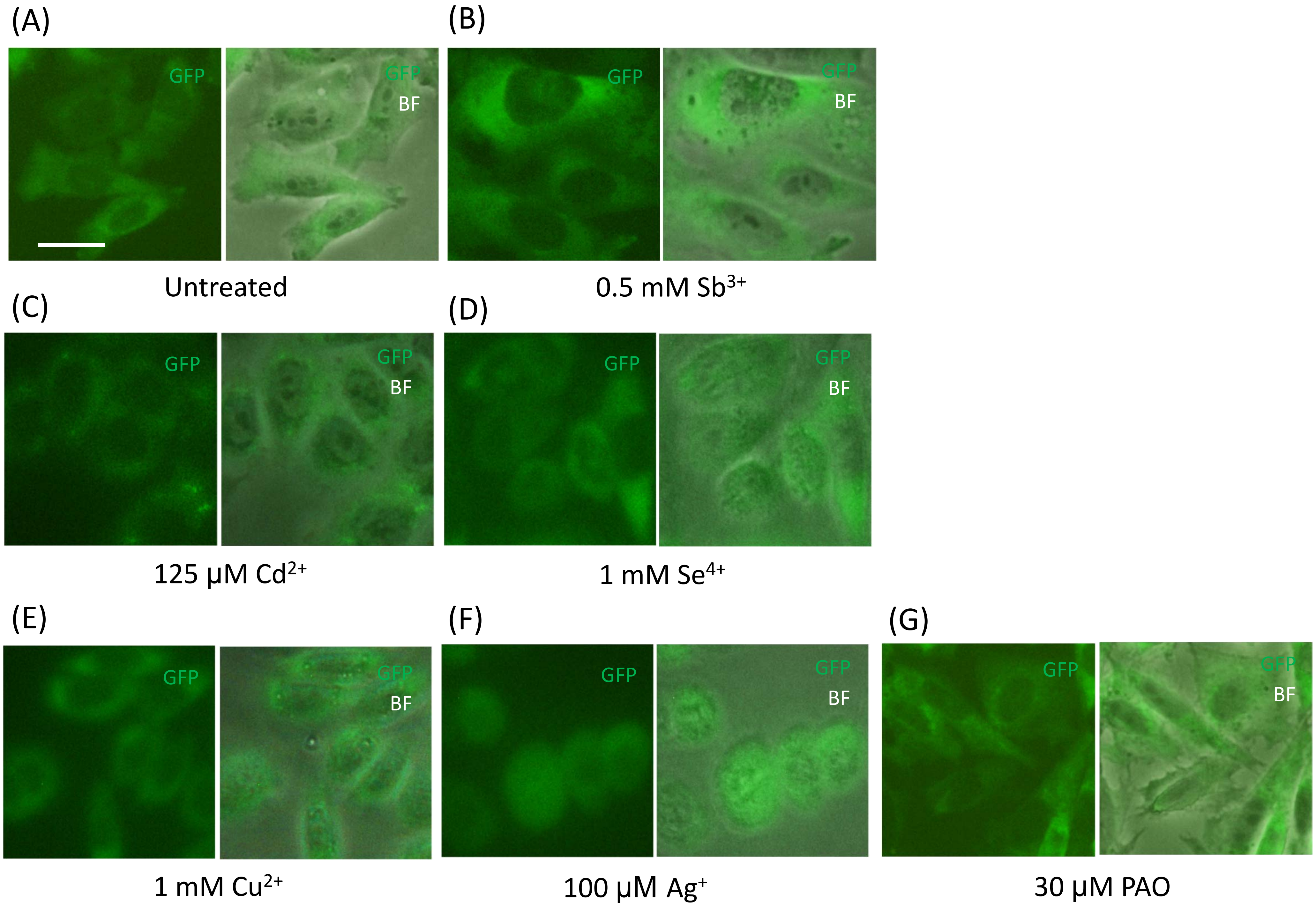


Supplementary Fig. 3

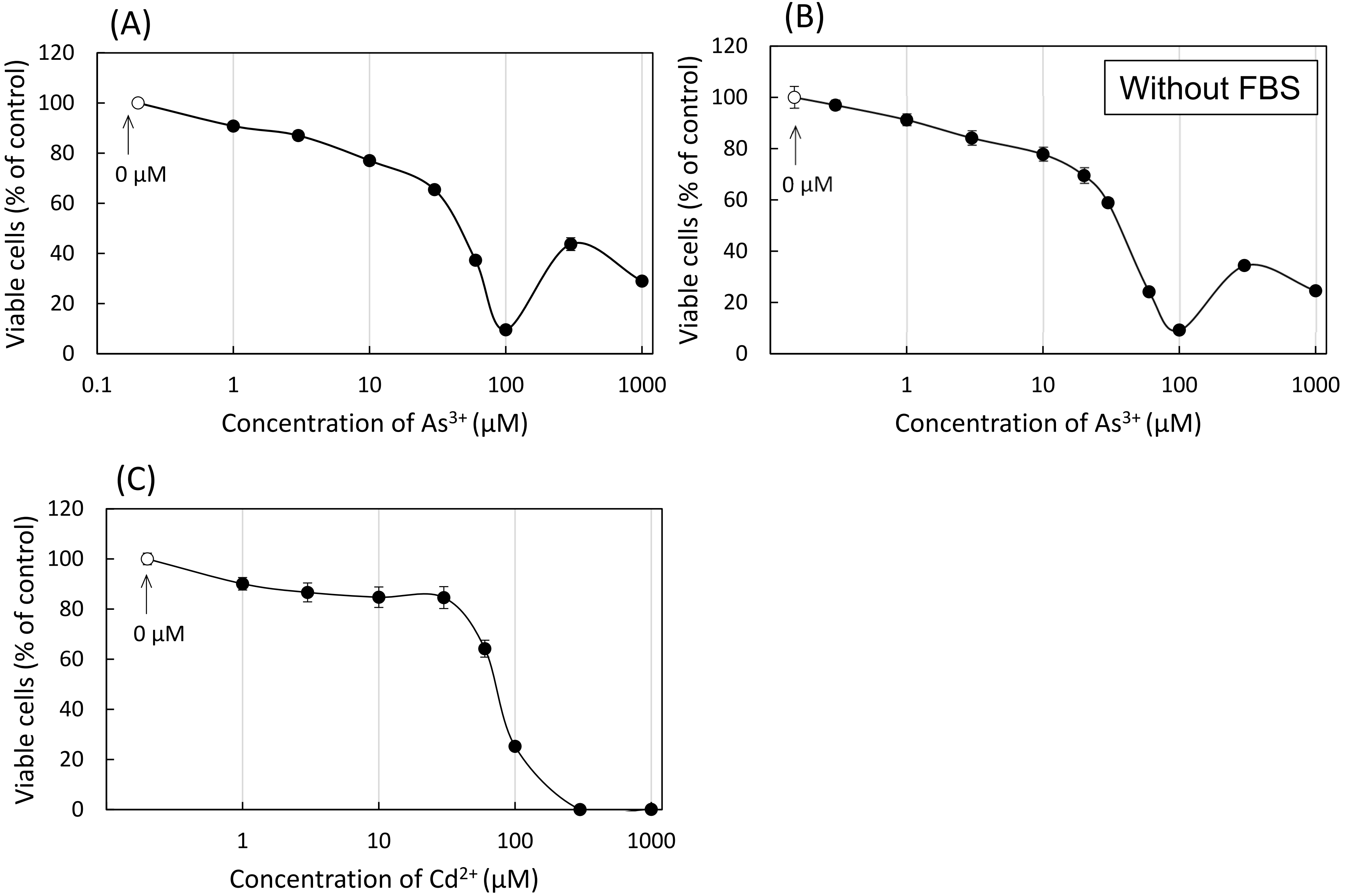
(A)
Viable cells (% of control)
0 µM
Concentration of As3+ (µM)
(B)
Without FBS
Viable cells (% of control)
0 µM
Concentration of As3+ (µM)
(C)
Viable cells (% of control)
0 µM
Concentration of Cd2+ (µM)

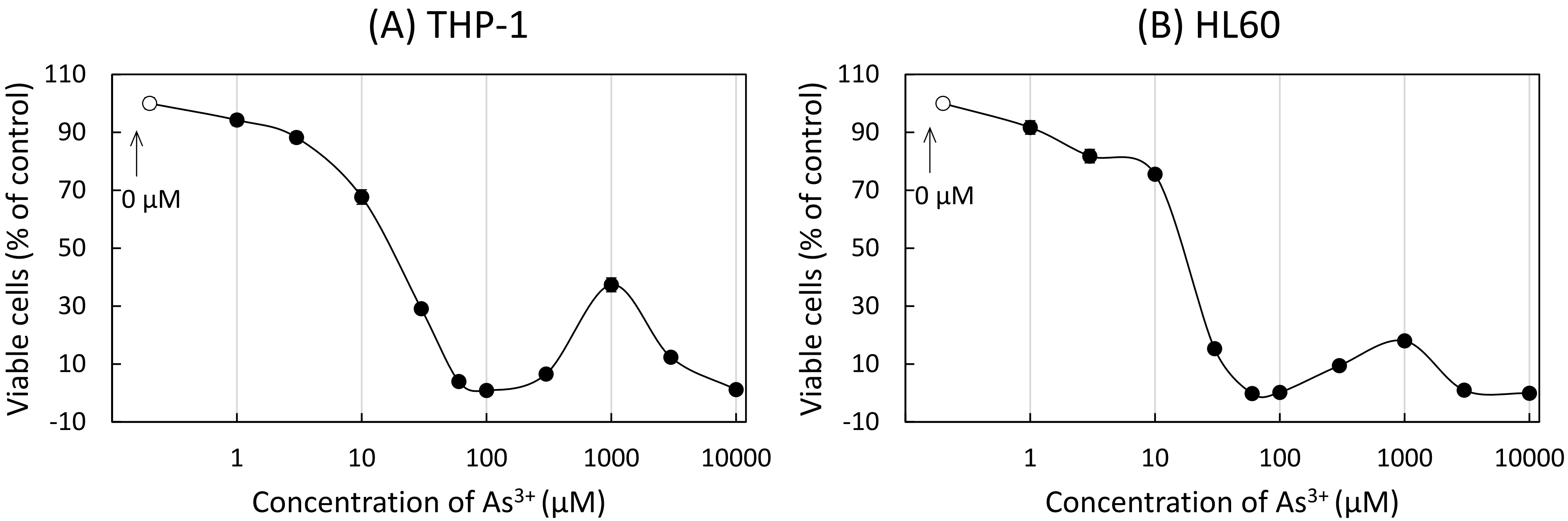


Supplementary Fig. 5

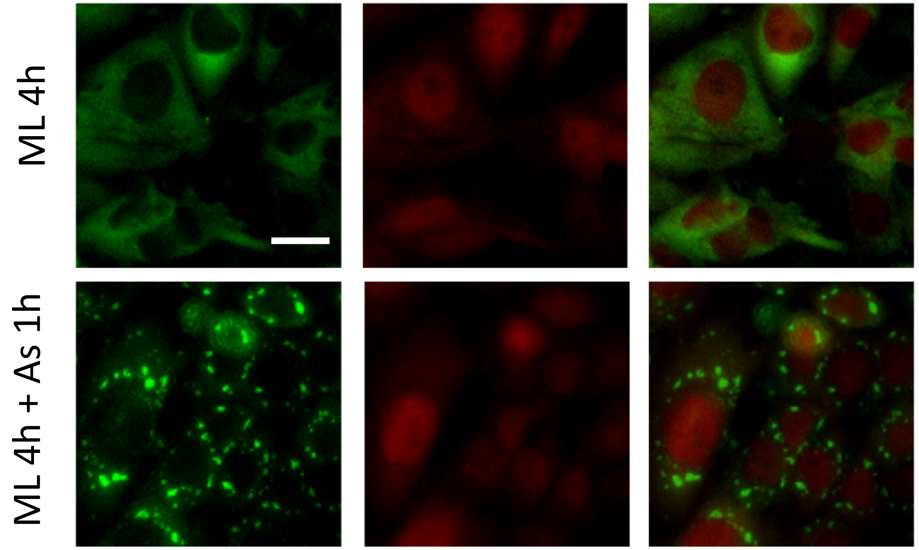


Supplementary Fig. 6
