## Supplementary material for "Formation of stress granules and non-canonical survival responses in arsenite-exposed cells": Source Table

Table 1, Information for antibodies and plasmids used in the present study.

| Reagent | Name | Source | Identifier | Dilution rate |
|---|---|---|---|---|
| Primary antibody | anti--eIF2α Mouse monoclonal | CST, Clone L57A5 | Cat# 2103 | 1:1000 for WB |
| | anti-phospho-eIF2α (Ser51) Rabbit monoclonal | CST, Clone D9G8 | Cat# 3398 | 1:1000 for WB |
| | anti-G3BP1, Mouse, monoclonal | Santa Cruz, Clone H-10 | Cat# sc-365338 | 1:1000 for WB |
| | anti-SUMO1 Rabbit, monoclonal | CST, Clone C9H1 | Cat# 4940 | 1:1000 for WB |
| | anti-SUMO2/3, Mouse, monoclonal, IgG2b | MBL, Clone 1E7 | Cat# M114-3 | 1:1000 for WB |
| | anti-PML, Rabbit, polyclonal | Bethyl | Cat# A301-167A-3 | 1:1000 for WB |
| | anti-ubiquitin Mouse, monoclonal | Cytoskelton, Clone P4D1 | Cat#: AUB01-S | 1:1000 for WB |
| | anti-GFP, Goat, polyclonal | Abcam | Cat# ab5450 | 1:1000 for WB |
| Secondary antibody | goat anti-Rabbit IgG (HRP) | Santa Cruz | Cat#: sc2054 | 1:2500 for WB |
| | goat anti-Mouse IgG (HRP) | Santa Cruz | Cat#: sc2055 | 1:2500 for WB |
| | mouse anti-Goat IgG (HRP) | Santa Cruz | Cat#: sc2354 | 1:2500 for WB |
| Conjugated antibody | anti-α-tubulin (HRP) Rabbit, polyclonal | MBL | Cat# PM054-7 Anti-α-tubulin HRP-DirectT | 1:2000 for WB |
| | anti-histone H3 (HRP) Rabbit, monoclonal, IgG2b | CST, Clone 3H1 | Cat# 12648 HRP-conjugated anti-histone H3 | 1:1000 for WB |
| Plasmid | GFP-tagged human G3BP1 transcript variant 1 | OriGene | RG202692 [NM_005754] | |
| | mCherry-SUMO2 Wild | VectorBuilder, mCherry/mSumo2 | VB210203-1053ufy [NM_133354.2] | |
| | mCherry-SUMO2 C-terminus GG deleted mutant | VectorBuilder, mCherry/mSumo2 | VB210203-1259pus [NM_133354.2] | |
