## Supplementary material for "Formation of stress granules and non-canonical survival responses in arsenite-exposed cells": Uncropped western image (Fig.7)

Fig. 7(A) Phospho-eIF2alpha

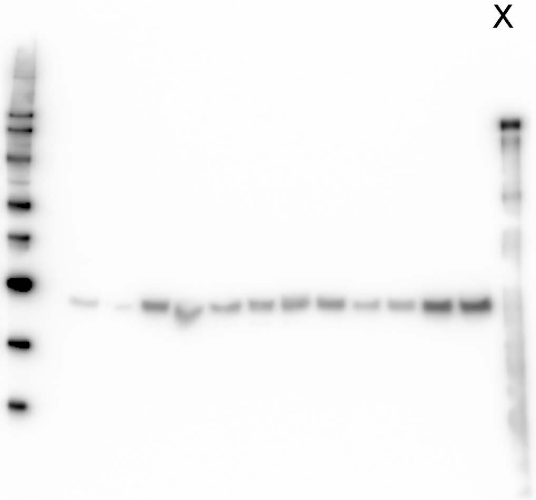

Fig.7(A) eIF2alpha

X

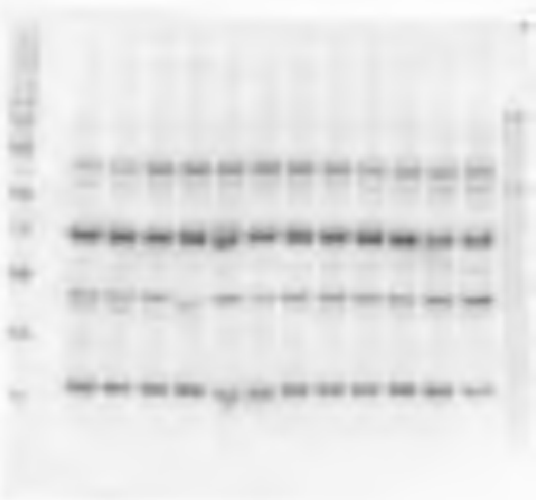

Fig.7(A) tubulin

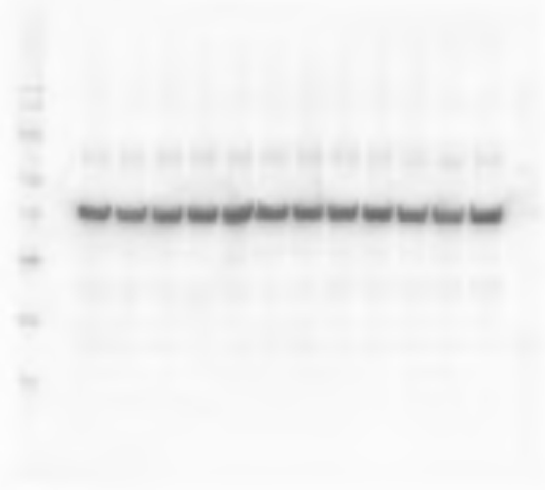

Fig.7(B) Phospho-eIF2alpha

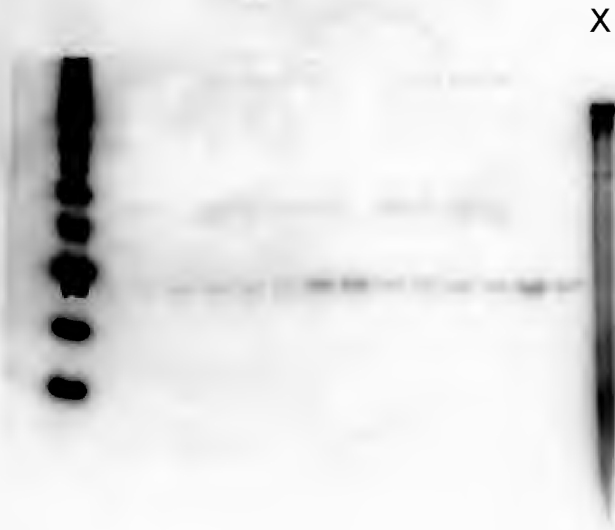

Fig.7(B) eIF2alpha

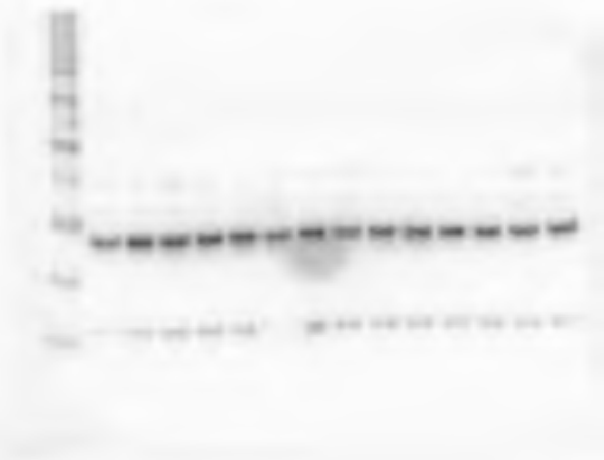

Fig.7(B) GFP

X

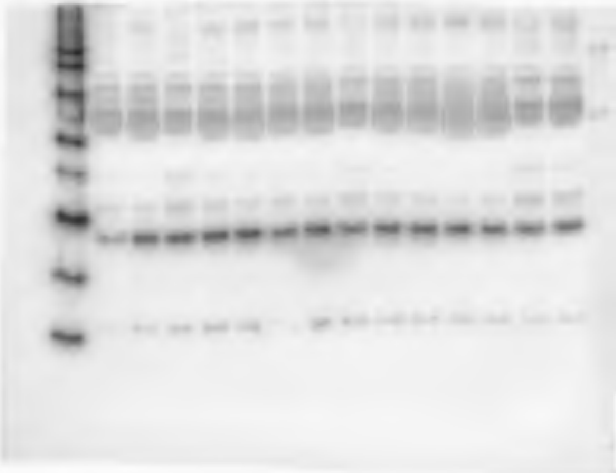
