## Supplementary material for "Formation of stress granules and non-canonical survival responses in arsenite-exposed cells": Uncropped western image (Fig.9)

FIg.9(B) PML

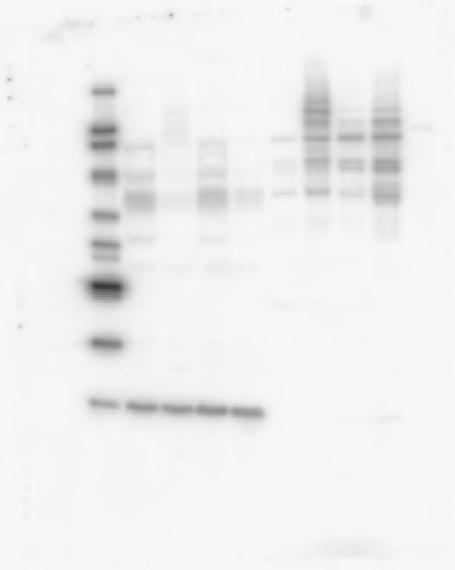

Fig.9(B) SUMO2/3

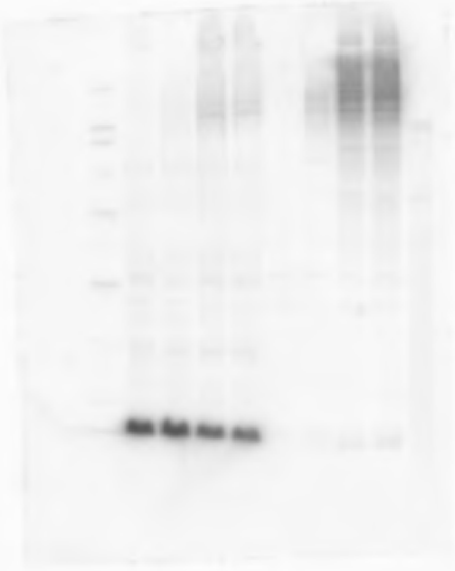

Fig.9(B) Tubulin + Hiatone H3 (Left)

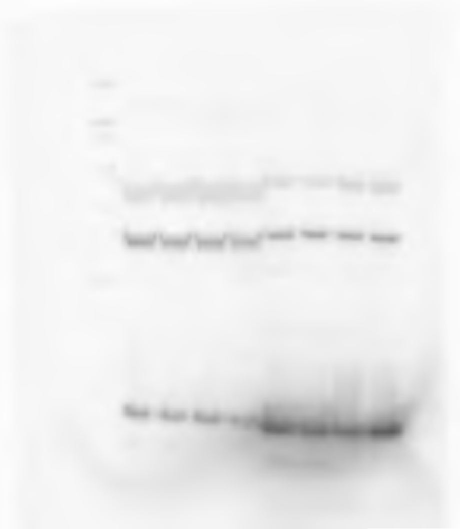

Fig.9(B) SUMO1

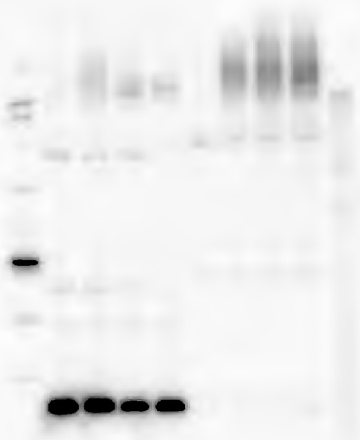

Fig.9(B) Ubiquitin

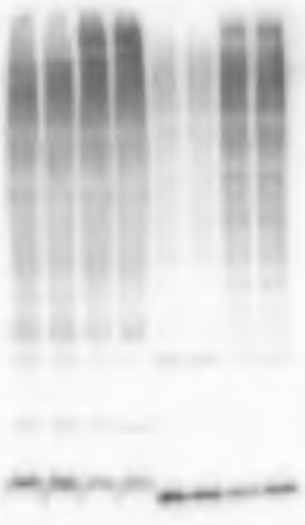

Fig.9(B) Tubulin + Hiatone H3 (Right)

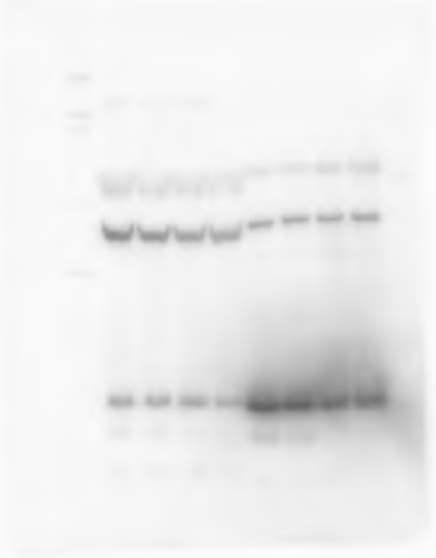
